# Fin elaboration via anterior-posterior regulation by Hedgehog signaling in teleosts

**DOI:** 10.1101/2023.10.10.557878

**Authors:** Yoshitaka Tanaka, Shun Okayama, Satoshi Ansai, Gembu Abe, Koji Tamura

## Abstract

Fins in fishes are appendages that serve to facilitate maneuvering in water. Compared to their ancestral state, teleosts have reduced radial bones in their paired fin skeletons and have acquired elaborated, agile paired fins. We found that mutation of *Hedgehog interacting protein* (*Hhip*), encoding an antagonist of Hedgehog signaling, leads to an increase of radial bones in a localized region and replicates the ancestral state. Interestingly, the caudal fin, which has undergone a reduction of skeleton structure in teleosts, as well as the paired fins, exhibit a regional-specific branching of the 2nd hypural in *hhip^-/-^* mutant zebrafish. These results imply that Hedgehog signaling is repressed during fin development in teleosts compared to the ancestral state. Molecular genetic analyses show that *shhb*, one of the *Sonic hedgehog* (*Shh*) genes, encoding one of the Hedgehog ligands in teleosts, is expressed during subdivision of endochondral components in paired fin skeletal development, and that mutation of *shhb* leads to hypoplasia of the paired fin skeletons. Therefore, we suggest that paired and caudal fins in fishes possess a specific region susceptible to Hedgehog signaling. The reduction of radial bones by repressive regulation of Hedgehog signaling may induce fin elaboration in teleosts.

## Introduction

Fins in fishes are the product of evolutionary adaptation and facilitate diverse hydrodynamic maneuvering, including locomotion, directional changes, and deceleration. It is particularly noteworthy that paired fins (pectoral and pelvic fins) have undergone morphological transformations resulting in a transition from their ancestral state to a derived form within the skeletal framework. In the ancestral state, pectoral fins possess a large number of radial bones as observed in chondrichthyans, some basal actinopterygians, placoderms, and acanthodians (1–6), and the pectoral fins of these fishes are composed of multiple basal radials, comprising the tribasal bones (propterygium, mesopterygium, and metapterygium) (5–7). On the other hand, pectoral fins in teleosts, derived fishes in actinopterygians, have lost these ancestral characteristics and have only a limited number of radials remaining (8–10). As a consequence of fin elaboration, teleosts utilize oscillatory motions of pectoral fins for propulsion, hovering, directional changes, or deceleration (11–14).

Fin elaboration by reduction of the number of radials has occurred along the anterior-posterior (AP) axis. In the development of vertebrate appendages, Hedgehog signaling regulates skeletal morphology along the AP axis through promotion of cell proliferation and patterning/differentiation of the skeleton. *Sonic hedgehog* (*Shh*) is one of the genes encoding a Hedgehog signaling ligand, and it regulates the number of skeletal components along the AP axis in the paired appendages (15). In limbs, which are paired appendages in tetrapods (amphibians and amniotes), and homologous to paired fins, the disruption or mutation of the limb-specific *Shh* enhancer results in hypo- or hyperplasia of digits along the AP axis (16–18). In paired fins, the disruption of *Shh* expression also produces severe defects in paired fin skeletons (19, 20). Therefore, it is plausible that *Shh* regulation along the AP axis is also implicated in the morphological evolution of the skeletons of paired fins in fishes.

Based on these previous reports, we hypothesize that Shh signaling, which is responsible for the induction of radials, in paired fins of teleosts is moderately repressed, thereby reducing the number of radials in them. To examine whether repression of Shh signaling affects the number of radials, we focused on the repressive mediators of the Hedgehog signaling pathway. We produced knockout models of repressive mediators of Hedgehog signaling in zebrafish and observed their development and alteration of paired fin skeletons.

## Results

### Hedgehog signaling overactivation by *hhip* mutations induces expansion of endochondral components in fin skeletons

As candidate genes for repressing Shh signaling, we examined two Hedgehog signaling repressive mediator genes, *Hedgehog interacting protein* (*Hhip*) and *GLI family zinc finger 3* (*Gli3*). *Hhip* mutant zebrafish are known to exhibit an increase in the number of fin rays and an expansion of pectoral fin primordia, but endochondral components in paired fin skeletons are not described (21, 22). *Gli3* mutant medaka are reported to exhibit an increase in the number of paired fin radials, but the fin radial phenotype in *Gli3* mutant zebrafish has not been reported (23). Using the CRISPR-Cas9 genome editing approach in zebrafish, we introduced a mutation in exon 4 of *hhip* (Fig. S1A) and in exon 5 of *gli3* (Fig. S2A) to replicate mutants reported previously (22, 23), and we isolated two *hhip* mutants (*hhip^H215RfsX1^*, *hhip^F242SfsX8^*: Fig. S1A) and a *gli3* mutant with a 122-bp deletion on exon5 (*gli3^Δ^*: Fig. S2A). Consequently, *hhip^-/-^*mutants showed an expansion of fin skeletons (Fig. S1B-E), while *gli3^Δ/Δ^*mutants showed no fin radial abnormalities (Fig. S2B-E). In *hhip*^-/-^ mutants, the endochondral disc of pectoral fins was elongated anteriorly (Fig. S1B-E). Therefore, we focused on the fin skeletons of adult *hhip*^-/-^ mutants.

In the pectoral fins of *hhip^-/-^* mutants, the pectoral girdle position was shifted laterally compared to that of WT zebrafish, which was similar to the position in chondrichthyans and basal actinopterygians (Fig. 1A and B). The pectoral fin skeletons of *hhip^-/-^* mutants showed an increased number of radials compared to WT zebrafish (Fig. 1C and D, and Fig. S1F). The pelvic fins of *hhip^-/-^* mutants showed no superficial differences in endochondral skeletons from WT (Fig. 1E and F). The *hhip^-/-^* mutation also affected median fins (Fig. 1G-L): dorsal, anal, and caudal. The dorsal fin of *hhip^-/-^* mutants showed an increased number of radials compared to WT zebrafish (Fig. 1G and H, and Fig. S1H, S3A and B). In contrast, the anal fin skeleton was completely absent in *hhip^-/-^* mutants (Fig. 1I and J, and Fig. S3B and D). In the caudal fin of *hhip^-/-^* mutants, the number of hypurals, bones formed ventral to the ural centra and supporting caudal fin, increased compared to WT zebrafish (Fig. 1K and L). The number of the hypurals in the region ventral to the hypural diastema, the gap between the 2nd and 3rd hypurals (Fig. 1K and L, the black dotted line), increased from two to five (Fig. 1K). Taken together, *hhip^-/-^* zebrafish mutants exhibited an increase in radials or hypurals for some fin skeletons. Considering that early disruptions of *hhip^-/-^* mutants are rescued by Hedgehog-repressive treatment (22), these results imply that Hedgehog signaling overactivation by *hhip* mutations induced expansion of fin skeletons.

**Fig. 1.**
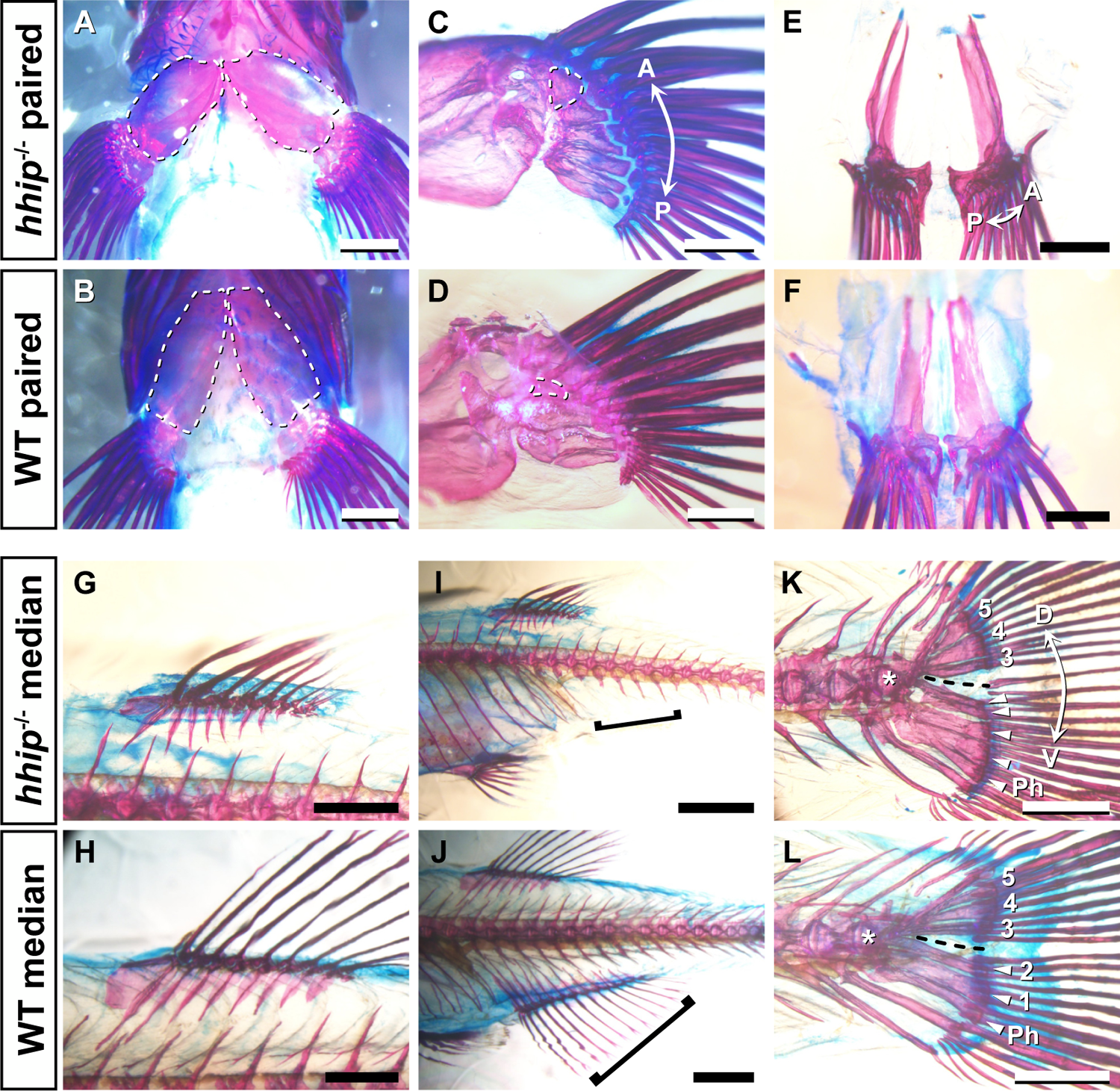
Paired and median fin skeletons in *hhip^-/-^* zebrafish. (A, B) Ventral view of pectoral fin skeleton of *hhip^-/-^* (A) and WT zebrafish (B). The white dotted lines show pectoral girdles. (C, D) Medial view of pectoral fin skeleton of *hhip^-/-^* (C) and WT zebrafish (D). The white dashed lines show the 1st proximal radial (PR1). (E, F) Pelvic fin skeleton of *hhip^-/-^*(E) and WT zebrafish (F). (G, H) Dorsal fin skeleton of *hhip^-/-^*(G) and WT zebrafish (H). (I, J) Anal fin skeleton of *hhip^-/-^* (G) and WT zebrafish (H). The black bracket indicates the post-anal region where the anal fin is formed. (K, L) Caudal fin skeleton of *hhip^-/-^* (K) and WT zebrafish (L). The black dashed line shows the hypural diastema, and the white arrowheads show hypurals in the region ventral to the hypural diastema. Numbers indicate the 1st to 5th hypurals. The white asterisk marks the 1st ural vertebra. Ph, parhypural. All observations were performed on 5 *hhip^-/-^* and 5 WT fish. Double arrows indicate the anterior (A)-posterior (P) axis and the dorsal (D)-ventral (V) axis. Scale bars: 500 µm (C-F); 1 mm (A, B, G, H, K, L); 2 mm (I, J).

### Regional-specific expansion in paired and caudal fin skeletons

Fishes that retain the ancestral state of paired fins show an increase in radials along the AP axis during development (4, 24, 25). To identify the regionality of these increases, we examined fin skeletons along the AP axis during the development of *hhip^-/-^* mutants (Fig. 2).

**Fig. 2.**
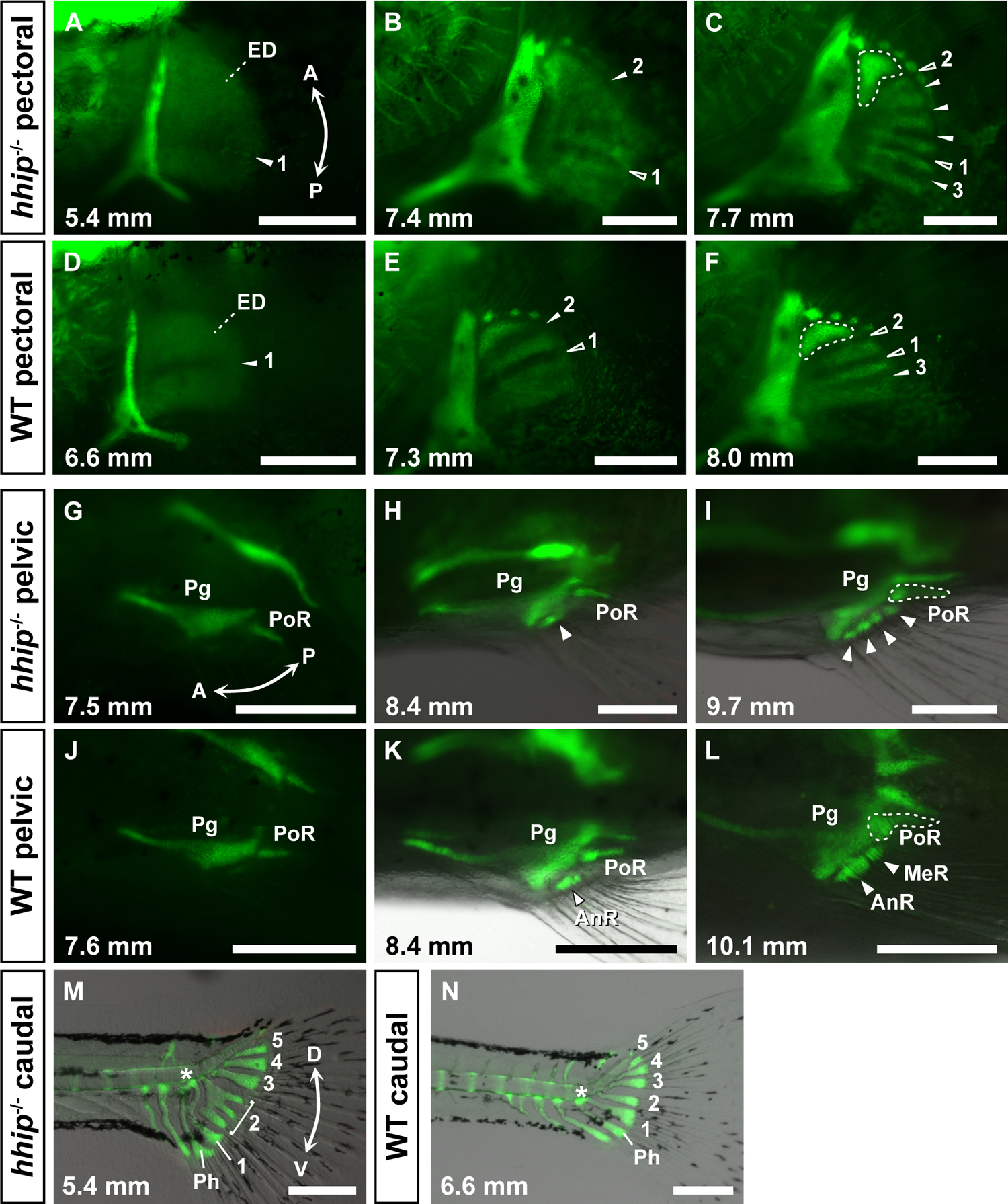
Developmental processes of the fin skeletons in *hhip^-/-^* zebrafish. (A-F) Pectoral fin development in *hhip^-/-^* (A-C) and WT zebrafish (D-F). Solid and open white arrowheads show new and previous cartilage subdivisions, respectively, and numbers mark the predicted 1st to 3rd subdivisions. White dashed lines show the 1st proximal radial (PR1). (G-L) Pelvic fin development in *hhip^-/-^* (G-I) and WT zebrafish (J-L). White arrowheads show the radial bones. AnR, anterior large radial; MeR, medial small radial; Pg, pelvic girdle; PoR, posterior elongated radial. (M, N) Caudal fin development in *hhip^-/-^* (M) and WT zebrafish (N). Numbers indicate the 1st to 5^th^hypurals. The white asterisk shows the 1st ural vertebra. Ph, parhypural. All observations were performed on more than 10 larvae at each developmental stage. Green fluorescence shows endochondral skeleton by *col2a1a:EGFP*. Double arrows indicate the anterior (A)-posterior (P) axis and the dorsal (D)-ventral (V) axis. Standard length (in mm) is given in the lower left of each panel. Scale bars: 250 µm (A-N).

In the development of WT pectoral fins, the 1st cartilage subdivision occurred in the middle of the endochondral disc, and the 2nd and 3rd cartilage subdivisions occurred in the anterior and posterior parts, respectively (9, 26) (Fig. 2D-F). In *hhip^-/-^*zebrafish, the 1st cartilage subdivision in *hhip* mutants shifted posteriorly (Fig. 2A, number 1), and the 2nd cartilage subdivision occurred in the anterior third (Fig. 2B, number 2). Subsequently, other cartilage subdivisions occurred (Fig. 2C, arrowheads without numbers). The posterior part further divided only once (Fig. 2C, number 3). Comparing the pectoral fin development of *hhip^-/-^*and WT zebrafish, the subdivision of the 1st proximal radial (PR1) in *hhip^-/-^*pectoral fins appears to be homologous to the 2nd cartilage subdivision in WT (Fig. 2E, number 2), and the subdivision on the posterior endochondral part appears to be homologous to the 3rd cartilage subdivision in WT (Fig. 2F). Comparing the subdivision sequence in *hhip^-/-^* mutants and WT, the position where proximal radials increased in the *hhip^-/-^*mutants appears to be in the anterior-median region, namely the 2nd proximal radial of the WT pectoral fins.

This developmental observation revealed an abnormality of the pelvic fin skeleton in *hhip^-/-^* mutants (Fig. 2G-L). In the development of WT pelvic fins, the skeletons possessed three radials (Fig. 2J-L): an anterior large radial (AnR), a medial small radial (MeR), and a posterior elongated radial (PoR). The anterior radials in *hhip^-/-^* mutants were as small as the MeR in WT but not as large as the AnR in WT (Fig. 2G-I). The most posterior radial was almost similar to the PoR in WT, suggesting that the *hhip* mutation did not affect the PoR (Fig. 2G-I). Based on these observations, the radials that increased in numbers in *hhip^-/-^* mutants were those in the anterior region of the WT pelvic fins (the AnR and MeR). In the development of the WT caudal fin, the parhypural and 1st hypural were formed ventral to the 1st ural vertebra (Fig. 2N, asterisk) and the 2nd hypural was generated ventral to the 2nd ural vertebra, respectively (27) (Fig. 2N). In *hhip^-/-^* mutants, the primordium of the 2nd hypural was branched and three hypurals were consequently formed (Fig. 2M), suggesting that hypurals increased in the anterior-median region of the caudal fin, which is occupied by the 2nd hypural in WT zebrafish. Therefore, in the paired and caudal fins of *hhip^-/-^*mutants, the expansion of radials or hypurals appears to be region-specific in the vicinity of the anterior-median region.

Next, to identify the AP regionality of fin skeletons in zebrafish from the perspective of molecular regulation, we visualized expression patterns of *hoxa/d13* genes, which were previously reported in some chondrichthyans and basal actinopterygians (28–30). Using the CRISPR-Cas9 knock-in approach in zebrafish (31–33), we introduced a EGFP reporter in *hoxa13b* and *hoxd13a* genes and made knock-in zebrafish lines (*hoxa13b^egfp^*, *hoxd13a^egfp^*: Fig. 3 and Fig. S4). Before the endochondral subdivision of pectoral fins, both *hoxa13b^egfp^*and *hoxd13a^egfp^* were expressed in the posterior region of the endochondral disc (Fig. 3A, B and D). During the endochondral subdivision of pectoral fins, *hoxa13b^egfp^* was expressed on the distal margin covering the 2nd, 3rd, and 4th proximal radials (Fig. 3C), and *hoxd13a^egfp^* was expressed on the distal margin covering the 3rd and 4th proximal radials (Fig. 3E). During pelvic fin development, while the *hoxa13b^egfp^* was expressed over the entire distal region (Fig. 3F-H), *hoxd13a^egfp^* was expressed in the posterior region where the PoR was formed (Fig. 3I and J). In the caudal fin primordium, *hoxa13b^egfp^* was expressed evenly until 2 dpf (Fig. 3K), and then median expression including the 2nd and 3rd hypurals was lost (Fig. 3L and M), while *hoxd13a^egfp^*was expressed only in blood vessels, not in the caudal fin primordium (Fig. 3N-P). Therefore, paired and caudal fins of zebrafish show regionality differentiated by *hoxa13* and/or *-d13* expressions.

**Fig. 3.**
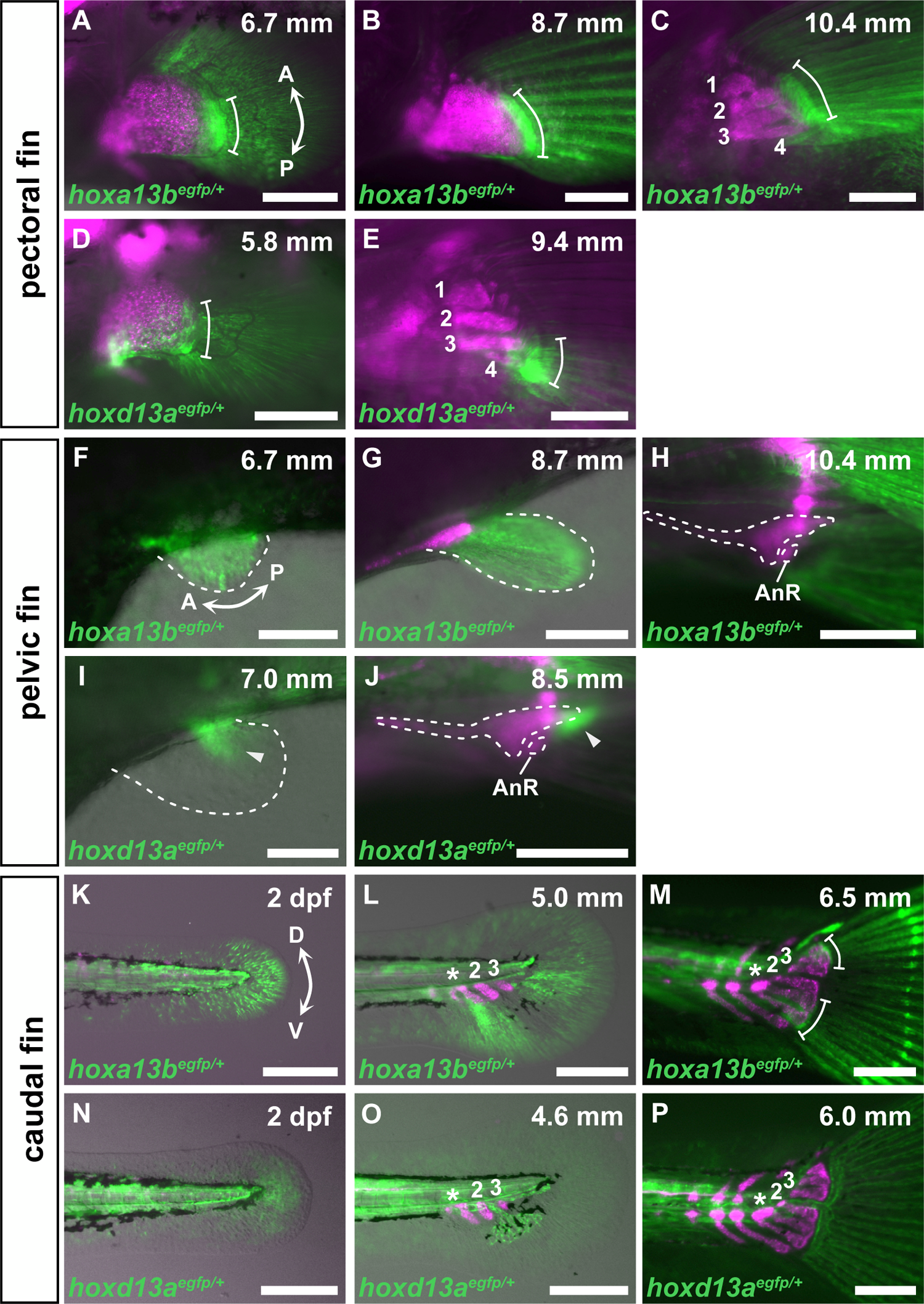
Expression of EGFP by *hoxa13b^egfp^* and *hoxd13a^egfp^*in paired and caudal fin development of zebrafish. (A-E) EGFP expression in pectoral fin development of *hoxa13b^egfp^* (A-C) and *hoxd13a^egfp^*(D, E). Numbers indicate the 1st to 4th proximal radials. White bars show the range of EGFP expression. (F-J) EGFP expression in pelvic fin development of *hoxa13b^egfp^* (F-H) and *hoxd13a^egfp^* (I, J). White arrowheads show EGFP expression. AnR, anterior large radial. (K-P) EGFP expression in caudal fin development of *hoxa13b^egfp^* (K-M) and *hoxd13a^egfp^* (N-P). Numbers indicate 2nd and 3rd hypurals. The white asterisk marks the 1st ural vertebra. White bars show the range of EGFP expression. All observations were performed on more than 10 larvae at each developmental stage. Magenta fluorescence shows paired fin skeleton by *sox10:DsRed*. Double arrows show the anterior (A)-posterior (P) axis and dorsal (D)-ventral (V) axis. Standard length (in mm) is given in the upper right of each panel. Scale bars: 125 µm (F, I); 250 µm (A-E, G, H, J-P).

### *Shh* serves as Hedgehog signaling ligands during late development of paired fins in teleosts

The increased expression of *hoxa13b* and *hoxd13a* with increasing proximity to the posterior region suggests AP regionality in developing radials. Given the subdivision process and the *hoxa13b* and *hoxd13a* expression pattern, region-specific expansion along the AP axis in *hhip^-/-^*mutants is regulated by the AP regionality derived from Hedgehog signaling. Hedgehog signaling in paired appendages is known to be related to these genes; signaling regulates expression of *hoxa13* and *hoxd13* in zebrafish (34) and is repressed by antagonization of Hhip (22, 35). However, expression patterns of Shh, a good candidate of Hedgehog ligands in paired appendages, remain unclear in the late development of paired fins. Although zebrafish have two *shh* genes, *shha* and *shhb* (Fig. S5), neither have been reported to be expressed during late fin development when fin skeletons are formed (after 7 dpf in zebrafish). To investigate whether *shha* and *shhb* are expressed in the late development of zebrafish paired fins, we made the knock-in zebrafish lines *shha^egfp^* and *shhb^egfp^*(Fig. S6) and examined *shha* and *shhb* expression activities in zebrafish (Fig. 4A-N). Both *shha^egfp^* and *shhb^egfp^* were expressed in the posterior edge of the pectoral fins (Fig. 4A-H), but *shha^egfp^* expression appeared until 4 dpf (Fig. 4A) and disappeared by 7 dpf (Fig. 4B-D), while *shhb^egfp^* expression extended beyond 4 dpf (Fig. 4E-H). Thus, *shhb* was expressed for much longer than *shha* and was expressed until endoskeletal morphogenesis in the pectoral fins. In pelvic fin development, *shha^egfp^* expression in the posterior edge was quite short-term (Fig. 4I-K), while *shhb^egfp^* expression in the posterior edge extended until the beginning of pelvic girdle formation (Fig. 4L-N). Therefore, *shhb* is also mainly expressed in the late development stage of the pelvic fins. In contrast, both *shha^egfp^* and *shhb^egfp^* could not be detected in the median fin primordia (Fig. S7).

**Fig. 4.**
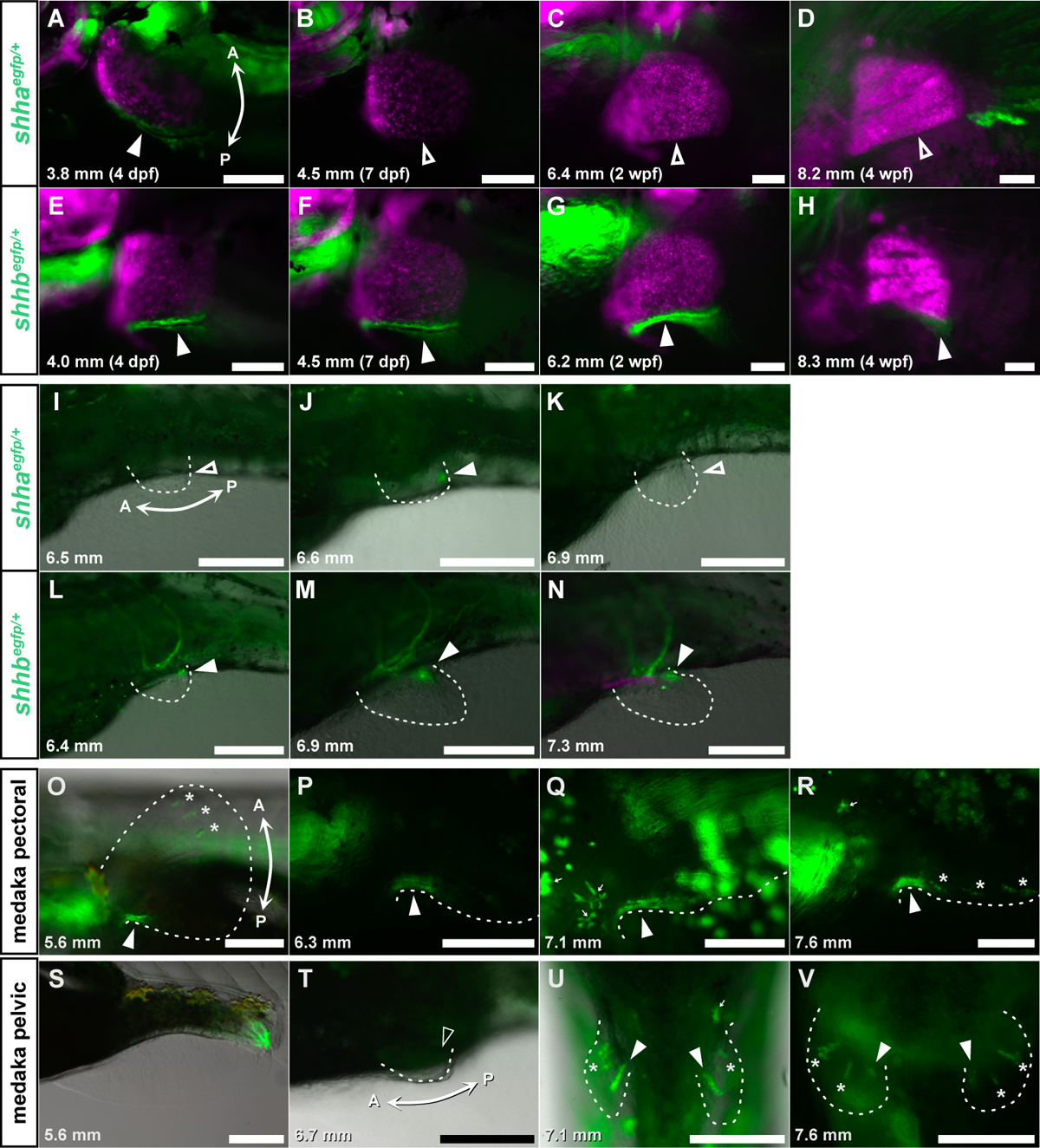
Expression of *shha^egfp^*and *shhb^egfp^* zebrafish, and *shha^egfp^* medaka in paired fin development. (A-H) EGFP expression in pectoral fin development of *shha^egfp^* (A-D) and *shhb^egfp^* zebrafish (E-H). (I-N) EGFP expression in pelvic fin development of *shha^egfp^*(I-K) and *shhb^egfp^* zebrafish (L-N). (O-R) EGFP expression in pectoral fin development of *shha^egfp^* medaka. (S-V) EGFP expression in pelvic fin development of *shha^egfp^* medaka. All observations were performed on 10 or more larvae at each developmental stage. Magenta fluorescence shows paired fin skeleton by *sox10:DsRed*. Solid white arrowheads show EGFP expression and open white arrowheads show no EGFP expression. Dashed lines show the outline of paired fin buds. The asterisk marks EGFP expression at the tips of fin rays. White arrows show autofluorescence by leucophores. Double arrows show the anterior (A)-posterior (P) axis. Standard length (in mm) is given in the lower left of each panel. Scale bars: 100 µm (A-H); 250 µm (I-V).

While basal teleosts including zebrafish have both *shha* and *shhb* genes, teleosts in Acanthomorpha including medaka, lost *shhb* and retained only *shha* (Fig. S5). If the *shha* expression pattern in medaka is similar to that in zebrafish, the teleosts in Acanthomorpha will have lost late developmental *Shh* expression corresponding to that of zebrafish after 7 dpf in paired fins. However, in most of the teleosts in Acanthomorpha, four radials are conserved in pectoral fin skeletons similar to basal teleosts which have both *shha* and *shhb* remaining (8). To explore how *shha* is expressed in Acanthomorpha, we made a knock-in medaka line (*shha^egfp^*: Fig. 4O-V and Fig. S8). The *shha^egfp^* medaka showed EGFP fluorescence in the posterior margin of developing pectoral fins (Fig. 4O-R). In the pectoral fins, *shha^egfp^* was expressed not only in the early stage (Fig. 4O and P) but also in the late stage when proximal radials began to be formed (Fig. 4Q and R). In pelvic fins, *shha^egfp^* was not expressed during the early budding (Fig. 4S and T) but was expressed from the mid-stage to the late stage (Fig. 4U and V). Similar to *shha^egfp^*and *shhb^egfp^* zebrafish, *shha^egfp^* medaka also showed no fluorescence in median fin primordia (Fig. S8B and C). Thus, the *shha* expression pattern in medaka was not similar to that of *shha* in zebrafish but to that of *shhb* in zebrafish. These data suggest that teleosts largely retain the *Shh* function in the late development of paired fins.

### Loss of *shhb* in zebrafish affects endochondral components in paired fin skeletons

To investigate the functions of late developmental *Shh* expression, we examined the function of *shhb* in zebrafish (Fig. 5). Using CRISPR-Cas9 genome editing in zebrafish, we introduced mutations in *shhb* and isolated one *shhb* mutant that was expected to encode a non-functional truncated protein (Fig. S9).

**Fig. 5.**
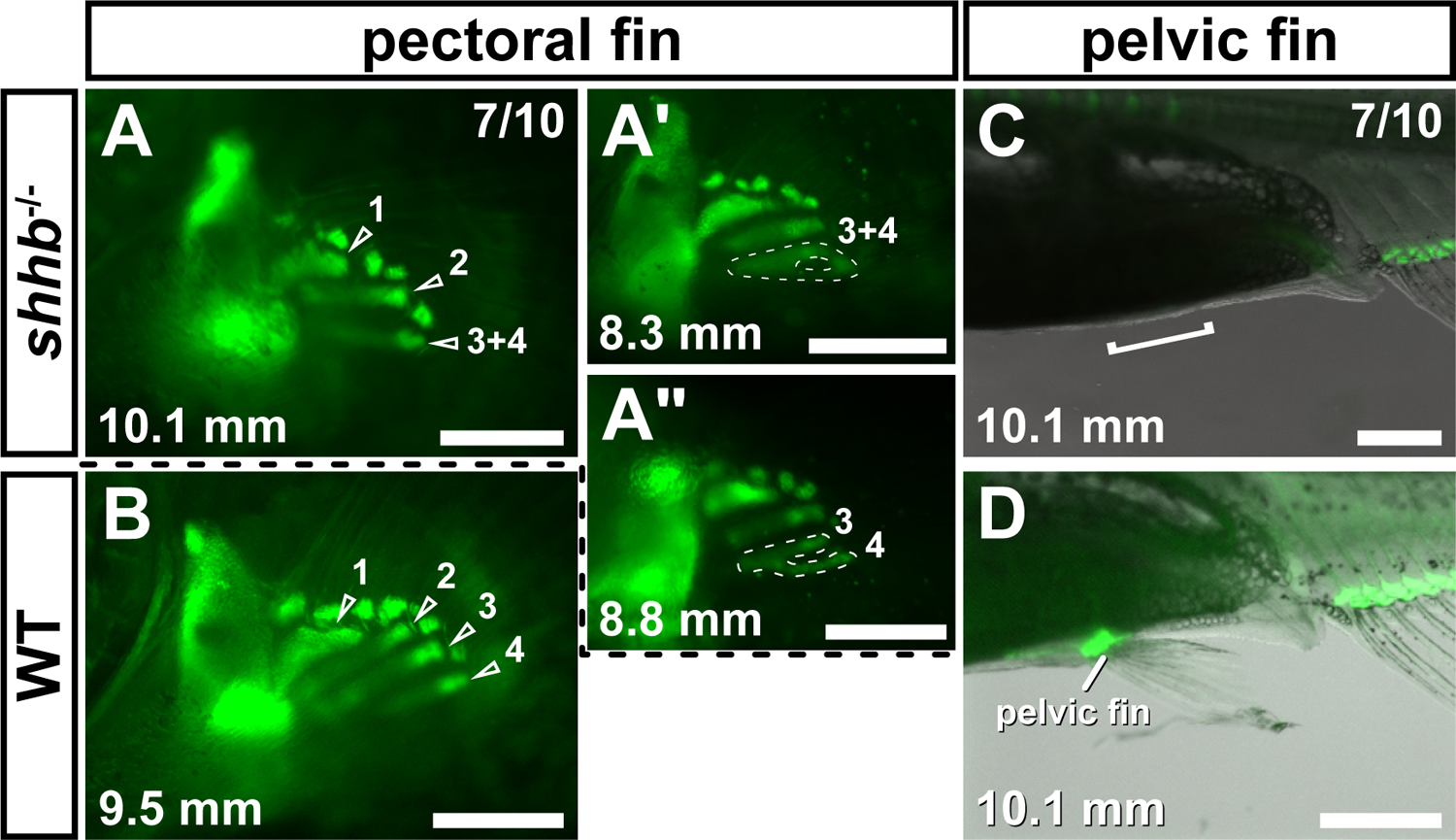
Paired fin skeletons in *shhb^-/-^* zebrafish. (A, B) Pectoral fin development in *shhb^-/-^* (A, n = 7/10) and WT zebrafish (B, n = 10/10). For *shhb^-/-^* mutants, 7/10 larvae exhibited complete fusion of the 3rd and 4th proximal radials in either left or right pectoral fins. A few *shhb^-/-^* zebrafish showed a mild defect of the subdivision between the 3rd and 4th proximal radials (A’, A’). Numbers indicate 1st-4th proximal radials. (C, D) Pelvic fin development in *shhb^-/-^* (C, n = 7/10) and WT zebrafish (D, n = 10/10). For *shhb^-/-^* mutants, 7/10 larvae showed pelvic fin loss, 6/10 larvae exhibited complete loss of both left and right pelvic fins, and 1/10 larvae exhibited complete loss of only the left pelvic fin. The white bracket indicates the pre-anal region where the pelvic fins are formed in WT zebrafish. Green fluorescence shows endochondral skeleton by *col2a1a:EGFP*. Standard length (in mm) is given in the lower right of each panel. Scale bars: 250 μm (A-A’, B); 500 μm (C, D).

In the pectoral fins of almost all *shhb^-/-^* mutant zebrafish, only three proximal radials were present (Fig. 5A). In some *shhb^-/-^* mutants, incomplete subdivision of the posterior endochondral part of the pectoral fin was observed, and the most posterior radial was fused 3rd and 4th proximal radials (Fig. 5A’ and A’). Therefore, late developmental *Shh* expression by *shhb* regulated the subdivision of the posterior endochondral part in pectoral fins. Surprisingly, pelvic fin skeleton, including pelvic girdles, was absent in almost all *shhb* mutants (Fig. 5C). In the development of zebrafish pelvic fins, *shha* was rarely expressed while *shhb* expression dominated (Fig. 4I-N). These results show that *shhb* constitutes the entirety of *Shh* expression and *shha* could not compensate for the absence of *shhb* in zebrafish pelvic fins. Thus, expression of *shhb* instead of *shha* was considered to regulate the entirety of the development of the pelvic fins.

## Discussion

We found that zebrafish *hhip^-/-^* mutation, in which Hedgehog signaling is overactivated, affects the fin skeleton, especially impacting the radials and hypurals. The phenotypes in *hhip^-/-^* mutants explain the developmental change in the morphological evolution from the ancestral form to the teleost-specific derived form. Considering the developmental process, the increase in radials in the pectoral fins of *hhip^-/-^*mutants occurred in the median region occupied by the 2nd proximal radial, not in the posterior region. In the pectoral fins of chondrichthyans and basal actinopterygians, the anterior (Fig. 6, red) and posterior basal radials (Fig. 6, green) are formed first, followed by the other median radials (Fig. 6, orange) which fill in the gap between the anterior and posterior radials (4, 24, 25, 36). Given the order in which radials are formed, the sequence of endochondral bone formation in the pectoral fin radials of some chondrichthyans and basal actinopterygians largely corresponds to that of *hhip^-/-^* mutant zebrafish (Fig. 2C). In addition, *Hox* gene expression supports the AP regionality based on endochondral subdivision during pectoral fin development in *hhip^-/-^* mutant zebrafish. *Hoxa13* is expressed on the distal margin of the median and posterior radials in the late development of paddlefish (28). *Hoxd13* is expressed in the posterior region of the pectoral fins in catshark and paddlefish (28–30). These *Hox* expression patterns correspond to those in zebrafish (Fig. 3A-J). Overall, whether *Hoxa13* and *Hoxd13* are expressed defines the anterior (neither *Hoxa13* nor *Hoxd13* are expressed), median (*Hoxa13* is expressed but *Hoxd13* is not expressed) and posterior (both *Hoxa13* and *Hoxd13* are expressed) regions in the pectoral fin skeleton. Given the increase of radials in *hhip^-/-^* mutants corresponding to the 2nd proximal radial (Fig. 3C and E, *Hoxa13* is expressed and *Hoxd13* is not expressed) in WT, the number of radials in the median region may be susceptible to Hedgehog signaling. Therefore, we suggest that the fin elaboration of pectoral fin skeleton along the AP axis occurs by the reduction of radials in the median Hedgehog-susceptible region (Fig. 6, orange) while the pectoral fins of teleosts are believed to have lost a posterior branching radial (the metapterygium) in the ancestral form, and have decreased to only four radials (6, 37). Pelvic fins of *hhip^-/-^* mutants also exhibit an increase in radials (Fig. 2I). The pelvic fin skeletons of chondrichthyans, sturgeon, and paddlefish possess multiple radials, while there are fewer radials in the pelvic fin skeletons of polypterus, bowfin, and teleosts (9, 24, 38–40). Therefore, Hedgehog signaling may also be involved in the variation of the number of radials in pelvic fins (Fig. S10).

**Fig. 6.**
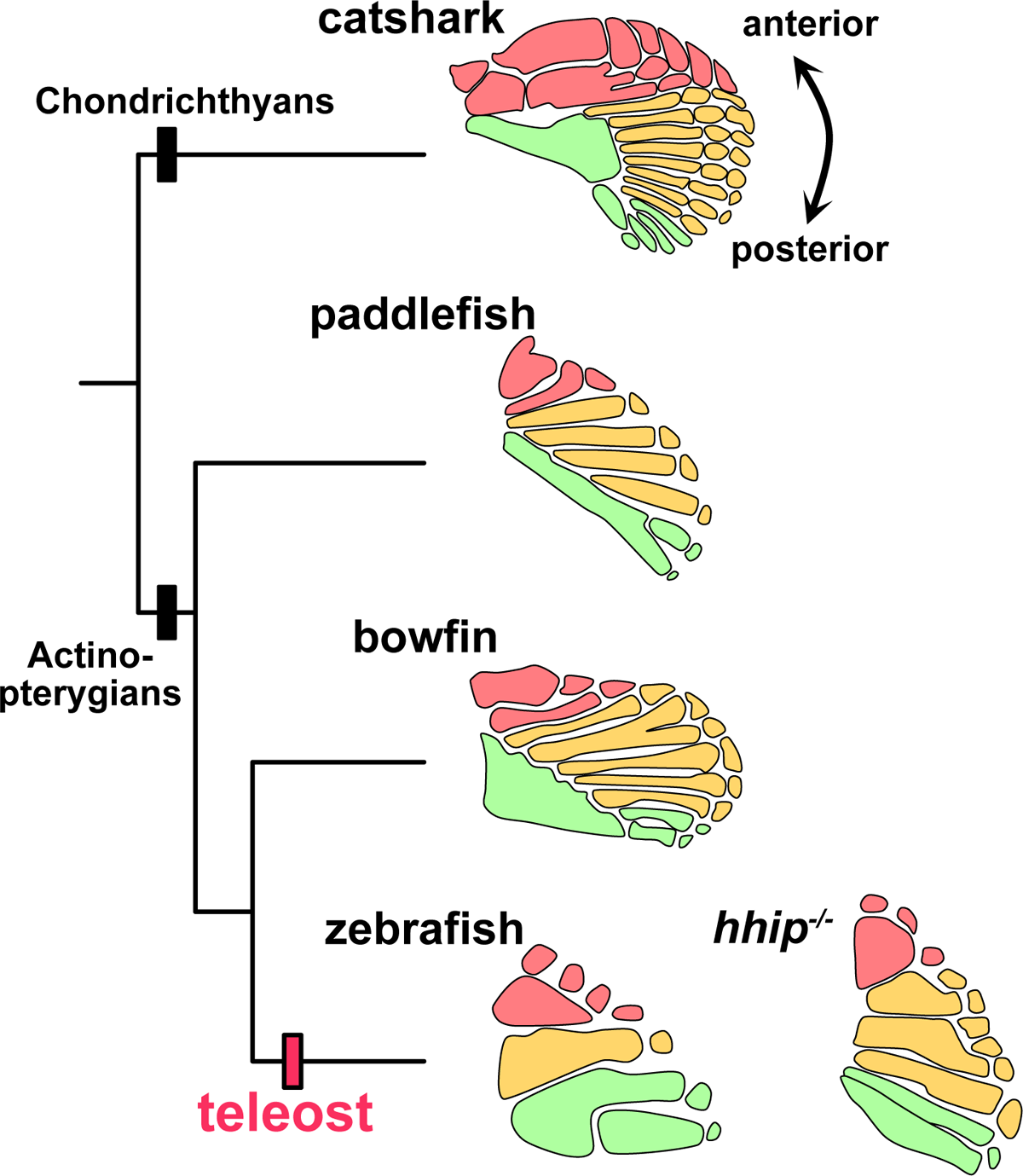
Elaboration of pectoral fins in teleosts. Colors indicate the anterior region where neither *Hoxa13* nor *Hoxd13* are expressed (red), the median region where *Hoxa13* is expressed but *Hoxd13* is not expressed (orange), and the posterior region where both *Hoxa13* and *Hoxd13* are expressed (green) based on the expression pattern of zebrafish and previous reports (28–30). The skeletons were redrawn with modification from the original sources as follows: catshark (5, 25), paddlefish (4, 24) and bowfin (37).

These patterns can be extended to caudal fin morphogenesis in teleosts. In the caudal fin skeleton of teleosts, the number of hypurals has decreased in parallel with paired fin elaboration (27). In *hhip^-/-^* mutants, hypural 2 was branched and three hypurals were formed (Fig. 2M), suggesting that caudal fin morphogenesis is regulated by axis formation similar to the AP regulation for paired fins. *Alx4*, an anterior-specific gene in the developing paired fins, is expressed at both the dorsal and ventral margins during caudal fin development, suggesting that the caudal fin possesses bipolar anterior regulation (41). In addition, the *gdf5* mutation in zebrafish, which affects the formation of posterior radials in pectoral fins, disturbs the formation of the dorsal hypural 3-5, suggesting that the dorsal region of the caudal fin may correspond to the posterior region of pectoral fins (42). These reports of molecular markers suggest that the dorsal-ventral axis in the caudal fin is regulated by a mechanism corresponding to the AP axis in paired fins (Fig. S10). Previous reports identified various phenotypes in the caudal fin skeleton (43, 44), but these results are the first report of Hedgehog-susceptible and regional-specific phenotypes. Although neither *shha* nor *shhb* expression were detected in the endochondral primordium of the caudal fin skeleton (Fig. S7A, B, E, and F), Shh secreted by other organizers, such as the notochord, regulate hypural development (45). We consider that Hedgehog susceptibility in the caudal fin may be involved in morphological evolution of the caudal fin skeleton. The number of hypurals decreased in the morphological change from the heterocercal form of chondrichthyans and basal actinopterygians to the homocercal form of teleosts (27), and this morphological change occurred approximately concurrently with paired fin elaboration. Therefore, Hedgehog signaling repression might affect not only paired fin evolution but also caudal fin evolution from the ancestral form to the teleost form.

Between basal teleosts (including zebrafish) and Acanthomorpha teleosts (including medaka), *Shh* regulation in paired fin development differs in several respects. The first discrepancy is that the repression mechanism of Hedgehog signaling differs. Our results showed that *hhip* mutant zebrafish exhibited an increase of radials but *gli3* mutant zebrafish exhibited no abnormality in the fin skeletons of zebrafish (Fig. S2). On the other hand, a previous report showed that *gli3* mutant medaka exhibit an increase of radials in the paired and dorsal fin skeletons (23). This difference may be related to the number of *Gli2* genes serving as GLI repressors (46, 47). For most teleosts, including zebrafish, two *Gli2* genes remain, but Japanese medaka (*Oryzias latipes*) has lost *gli2a* and only *gli2b* remains. Therefore, the loss of *gli3* function in Japanese medaka may produce severe effects compared to those in zebrafish due to weak compensation by only *gli2b*. The second discrepancy lies in in how *Shh* expression is regulated between zebrafish and medaka (Fig. 4). In zebrafish, both *shha* and *shhb* regulated paired fin development, while only *shha* regulated the development in medaka (Fig. 4). *Shh* expression at the posterior margin of paired fins is regulated by a single cis-regulatory element, known as the ZPA regulatory sequence (ZRS) (16, 17, 20). ZRS is located in intron 5 of *Lmbr1*, a gene on the same chromosome as *Shh*, and is common within gnathostome genomes. In basal teleosts, the genomic region where *shhb* is located has lost the *lmbr1* paralog and some neighboring genes and has become a kind of gene desert, while the genomic region where *shha* is located conserves the *lmbr1* paralog (Fig. S5). Considering that the expression pattern of *Shh* genes in paired fins of zebrafish and medaka, we speculate that zebrafish *shhb* remains a paired fin enhancer homologous to the ZRS of *shha* in the gene desert and achieves the fine-tuning regulation of *Shh* expression, while in medaka, the loss of one of the *Shh* genes simplifies such regulation. This diversity in Hedgehog signaling regulation may explain the morphological differences of fin skeletons within teleosts. In this respect, it is noteworthy that there are several morphological differences in the fin skeletons between basal teleosts and Acanthomorpha teleosts. For instance, pectoral fins in Acanthomorpha possess four hourglass radials that are uniform along the AP axis (8, 39, 48), similar to uniform and indistinguishable digits in limbs with the *gli3* mutation that disturbs the AP polarity (49, 50). The hourglass radials might reflect attenuation of AP regulation. Overall, we suggest that *Shh* regulation in paired fin development has changed between basal and Acanthomorpha teleosts and that this change has affected AP regulation which in turn leads to morphological differences within teleosts.

In summary, our findings showed fin elaboration via augmented repression of Hedgehog signaling and the diversity of *Shh* regulation within teleosts. However, it remains to be determined whether *Shh* regulates the AP identities of radials in teleosts. Further genetic analyses of the AP regulatory genes (e.g., *Alx4*, *Hand2*) or non-model species (e.g., chondrichthyans or basal actinopterygians) will reveal the precise evolutionary mechanism of fin skeletons in fishes.

## Supplementary figures

**Fig. S1.**
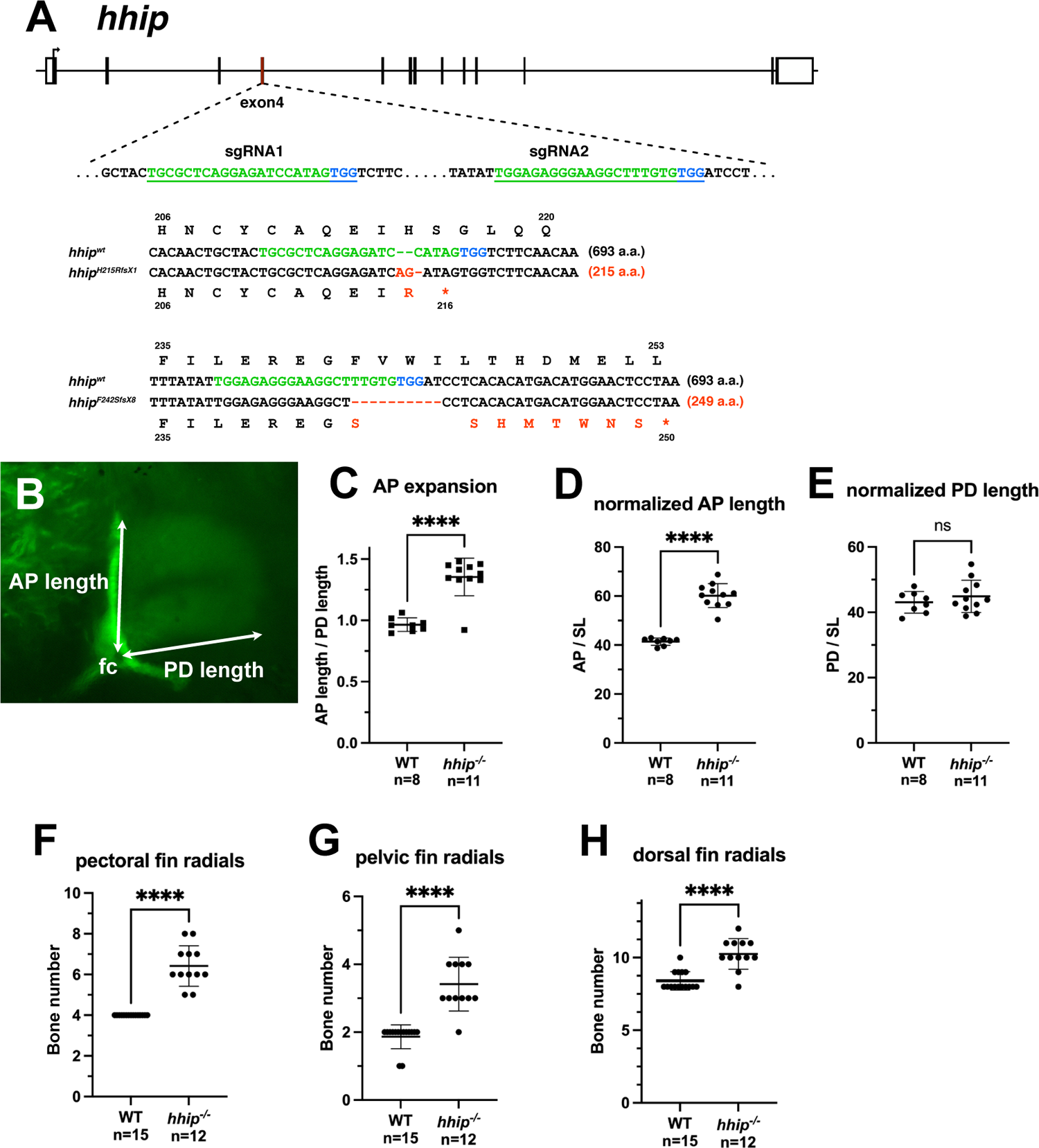
*hhip* mutation in zebrafish. (A) Sequence of *hhip* mutations. Blue sequences show the protospacer adjacent motif (PAM), green sequences show the recognition sequence for the sgRNA target and red sequences show the mutation site. (B) Measurement method of the length of pectoral fin skeleton. The anterior-posterior (AP) length was measured from foramen coracoideus (fc) to the anterior margin of the endochondral disc. The proximal-distal (PD) length was measured from foramen coracoideus to the distal margin of the endochondral disc. (C-E) Quantification of the length of pectoral fins in *hhip* mutants. Fish (SL = 5.0 – 7.5 mm) which did not start the subdivision of the endochondral disc were used in these analyses. An unpaired *t* test was used for the statistical analysis of pectoral fin length. (C) AP length normalized by PD length. Pectoral fins of *hhip^-/-^* mutants expanded along the AP axis. \*\*\*\**p*<0.0001, WT (mean 0.9651), *hhip^-/-^* (mean 1.354). (D, E) AP and PD lengths normalized by standard length (SL). The pectoral fins of *hhip^-/-^*mutants did not expand along the PD axis. (D) \*\*\*\**p*<0.0001, WT (mean 41.41), *hhip^-/-^* (mean 60.18). (E) *p*=0.3077, WT (mean 43.06), *hhip^-/-^* (mean 44.88). (F-H) Quantification of the number of radials in pectoral (F), pelvic (G) and dorsal fins (H). The number of pelvic fin radials did not include the PoR. Fish (SL > 8.0 mm) with complete formation of skeletal elements were used in these analyses. An unpaired *t* test was used for statistical analysis of skeletal element number. (F) \*\*\*\**p*<0.0001, WT (mean 4.00), *hhip^-/-^* (mean 6.417). (G) \*\*\*\**p*<0.0001, WT (mean 1.867), *hhip^-/-^* (mean 3.417). (H) \*\*\*\**p*<0.0001, WT (mean 8.40), *hhip^-/-^* (mean 10.25).

**Fig. S2.**
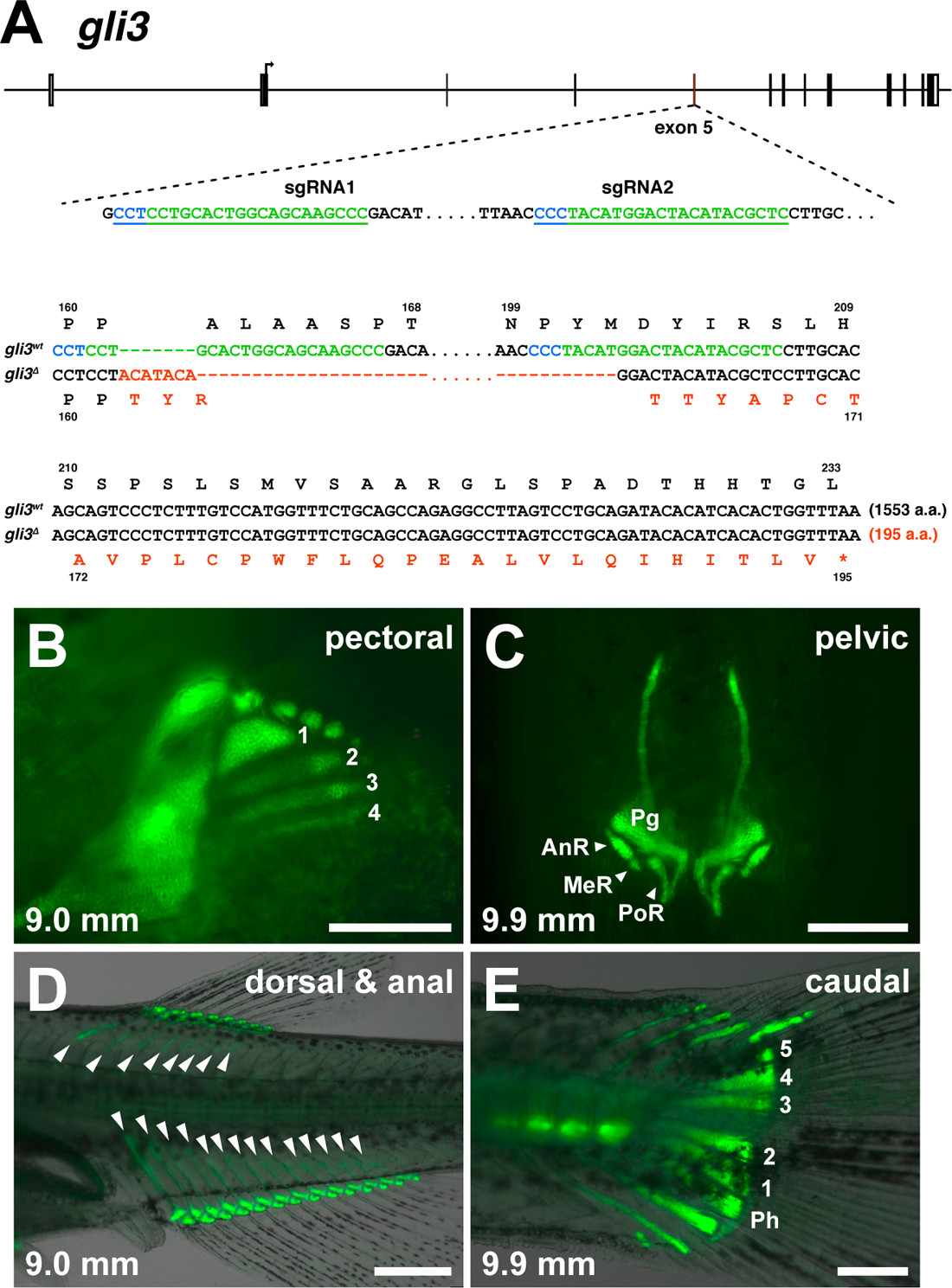
*gli3* mutation in zebrafish. (A) Sequence of *gli3* mutations. Blue sequences show the protospacer adjacent motif (PAM), green sequences show the recognition sequence for the sgRNA target, and red sequences show the mutation site. (B) Pectoral fin skeleton in *gli3^Δ/Δ^*. Numbers indicate the 1st to 4th proximal radials. (C) Pelvic fin skeleton in *gli3^Δ/Δ^*. AnR, anterior large radial; MeR, medial small radial; Pg, pelvic girdle; PoR, posterior elongated radial. (D) Dorsal and anal fin skeletons in *gli3^Δ/Δ^*. Closed white arrowheads show the radial bones. (E) Caudal fin skeleton in *gli3^Δ/Δ^*. Numbers indicate the 1st to 5th hypurals. Ph, parhypural. Observations were performed on 5 or more larvae with *gli3^Δ/Δ^* mutants. Standard length (in mm) is given in the lower left in panels (B-E). Scale bars: 250 µm (B, C, E); 500 µm (D).

**Fig. S3.**
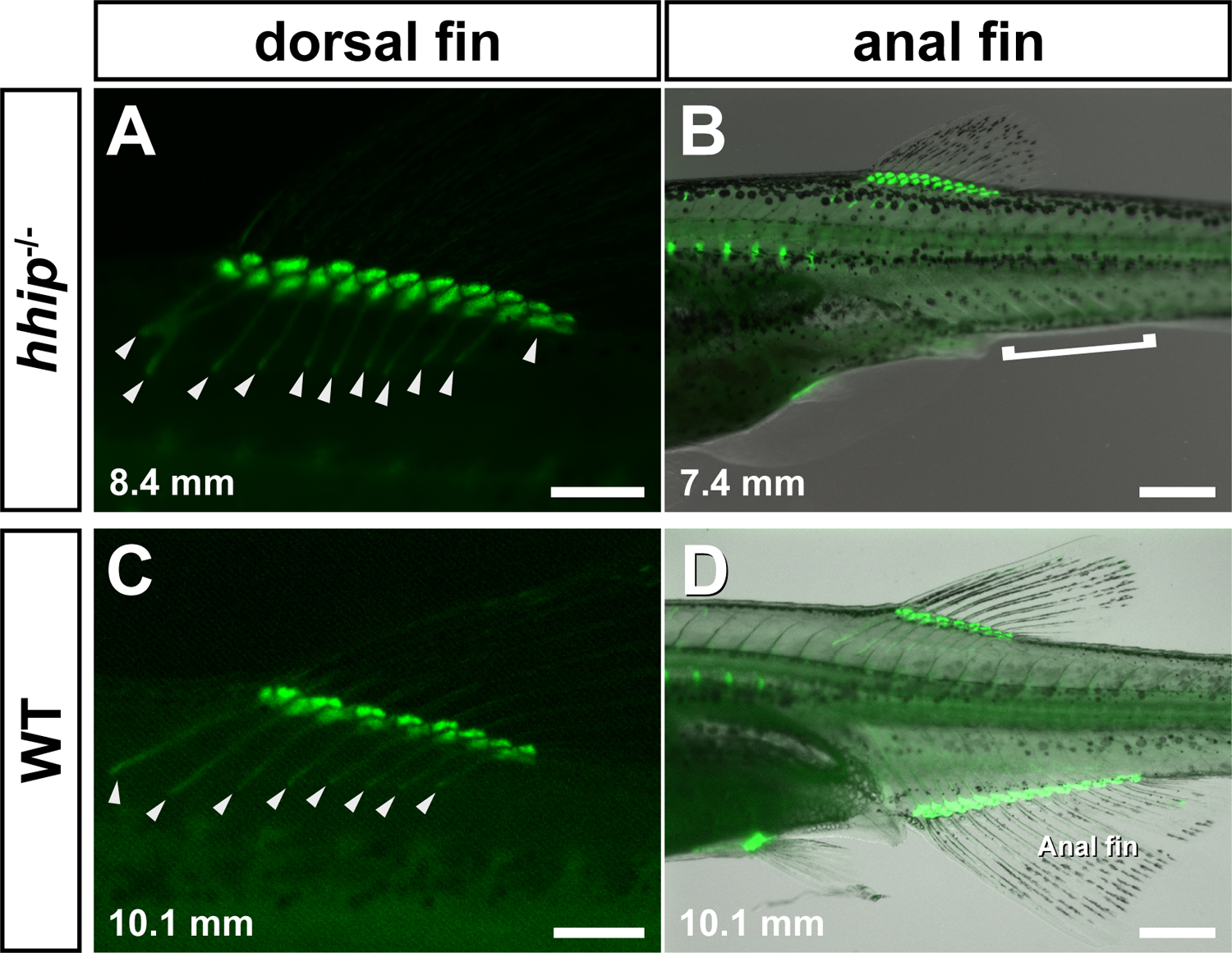
Dorsal and anal fin skeletons in *hhip* mutant zebrafish. (A, B) Dorsal and anal fin skeletons in *hhip^-/-^* zebrafish. (C, D) Dorsal and anal fin skeletons in WT zebrafish (D). Solid white arrowheads show the radial bones. The white bracket shows the post-anal region where the anal fin is formed in WT zebrafish. These observations were made on 10 or more *hhip^-/-^* and WT larvae. Green fluorescence shows endochondral skeleton by *col2a1a:EGFP*. Standard length (in mm) is given in the lower left of each panel. Scale bars: 250 µm (A, C); 500 µm (B, D).

**Fig. S4.**
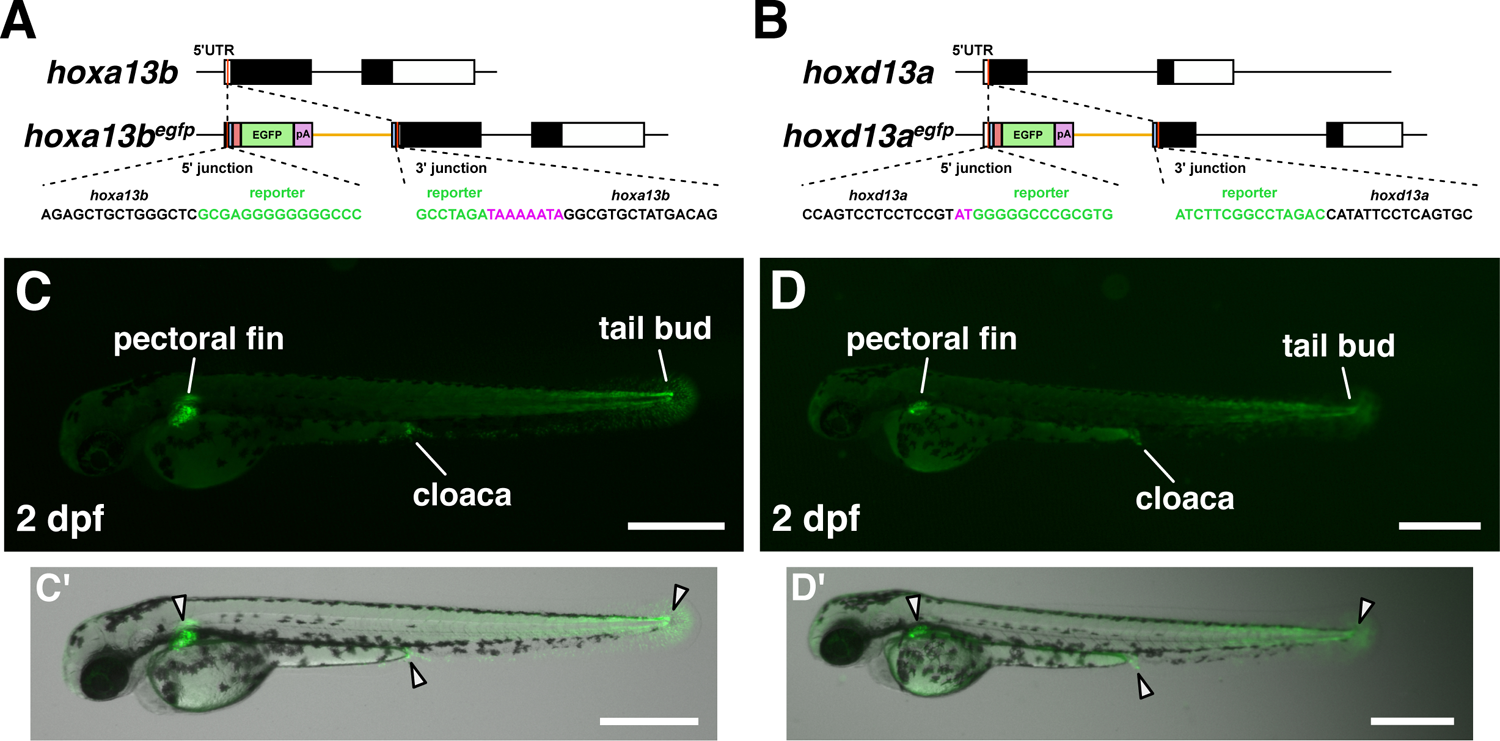
*hoxa13b^egfp^* and *hoxd13a^egfp^* knock-in zebrafish. (A, B) Sequences at the 5’ and 3’ junction of the *hoxa13b^egfp^* and *hoxd13a^egfp^*zebrafish. Magenta letters represent a junk insertion by CRISPR-Cas9 editing. (C, D) Overall views of EGFP expressions in 2 dpf zebrafish of *hoxa13b^egfp^* (C, C’) and *hoxd13a^egfp^* (D, D’). Solid white arrowheads show EGFP expression in the bright-field. These observations were made on 10 or more *hoxa13b^egfp^*and *hoxd13a^egfp^* zebrafish larvae. Scale bars: 500 µm (C, C’, D, D’).

**Fig. S5.**
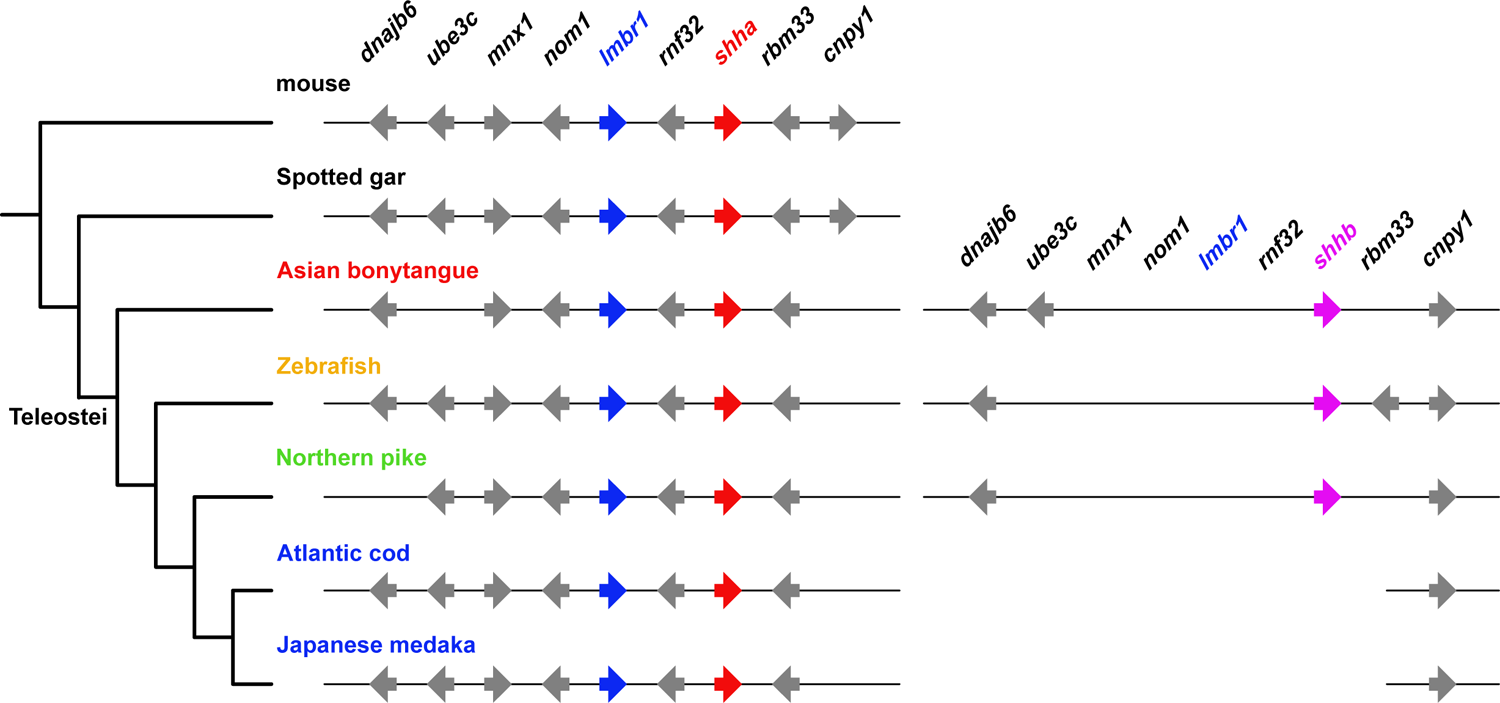
Comparison of *Shh* microsyntenic regions. Arrows show genes and the arrow directions show the 5’ to 3’ directions of the genes. The red arrows show *shha* (or *Shh* in vertebrates except teleosts) and the magenta arrows show *shhb*. The blue arrows show *Lmbr1*, which includes the ZPA regulatory sequence (ZRS) in the intron 5. Teleosts in Acanthomorpha have chromosomal recombination upstream of *cnpy1* and have lost *shhb* and the other microsyntenic genes.

**Fig. S6.**
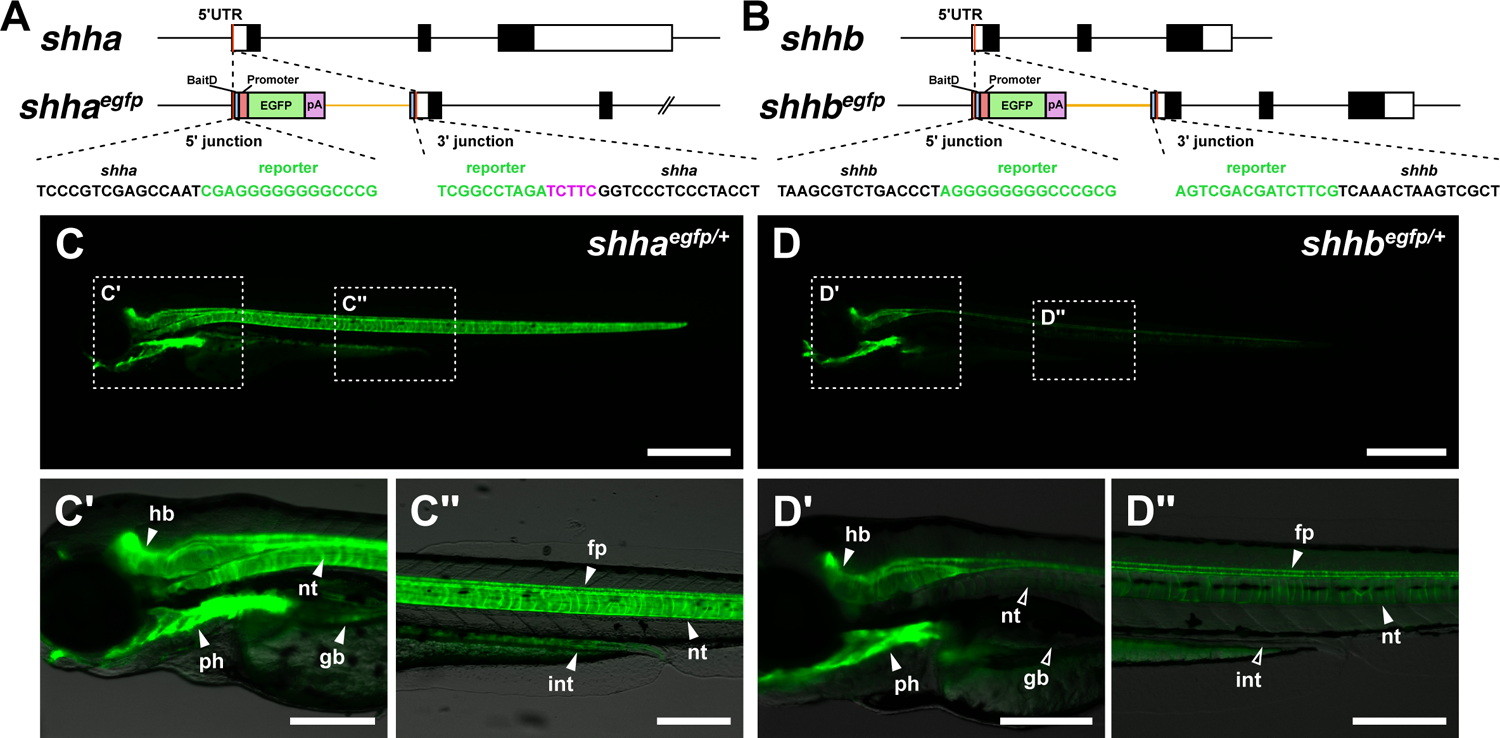
*shha^egfp^* and *shhb^egfp^* knock-in zebrafish. (A, B) Sequences at the 5’ and 3’ junction of *shha^egfp^* and *shhb^egfp^* zebrafish. Magenta letters represent a junk insertion by CRISPR-Cas9 editing. (C, D) Overall views of EGFP expressions in 3 dpf zebrafish of *shha^egfp^* (C) and *shhb^egfp^* (D). Solid white arrowheads indicate EGFP expression, and open white arrowheads indicate no EGFP expression. fp, floor plate of neural tube; gb, gas bladder; hd, hindbrain; int, intestine; nt, notochord; ph, pharynx. These observations were made on 10 or more *shha^egfp^* and *shhb^egfp^*zebrafish larvae. Scale bars: 500 µm (C, D); 250 µm (C’, C’’, D’, D’’).

**Fig. S7.**
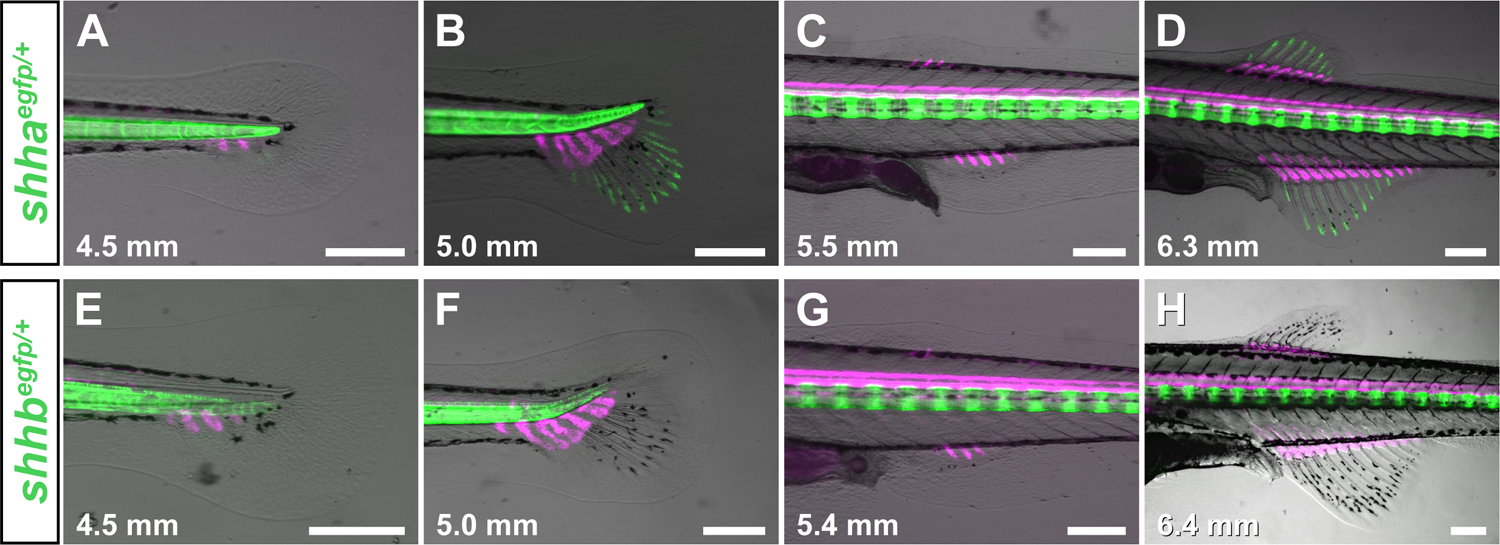
Expression of EGFP by *shha^egfp^* and *shhb^egfp^* in median fin development of zebrafish. (A-D) EGFP expression in median fin development of *shha^egfp^*. (A, B) Dorsal and anal fins. (C, D) Caudal fin. (E-H) EGFP expression in median fin development of *shhb^egfp^*. (E, F: Dorsal and anal fins. G, H: Caudal fin) Magenta fluorescence shows paired fin skeleton by *sox10:DsRed*. These observations were performed on 10 or more *shha^egfp^*and *shhb^egfp^* zebrafish larvae. Standard length (in mm) is given in the lower left of each panel. Scale bars: 250 µm (A-H).

**Fig. S8.**
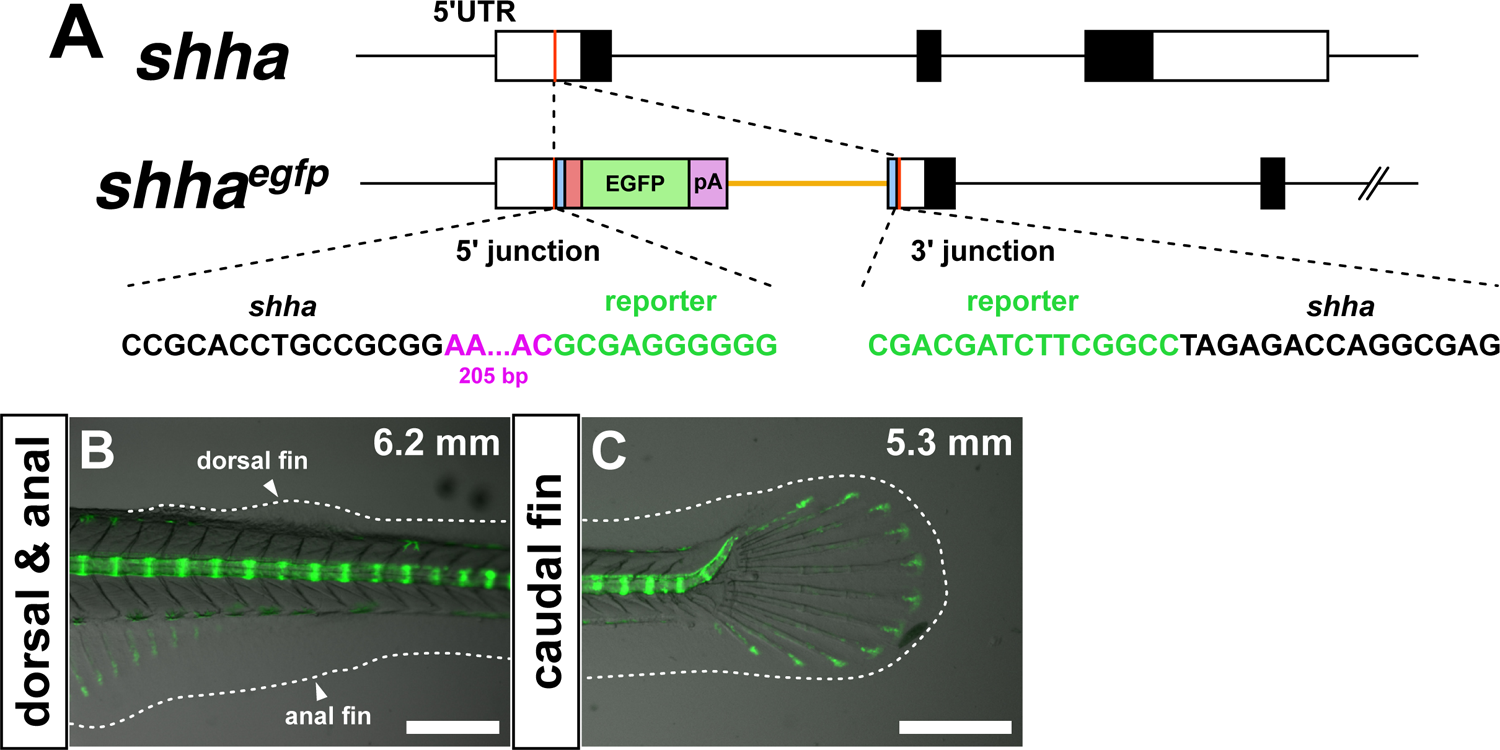
*shha^egfp^* knock-in medaka. (A) Sequences at the 5’ and 3’ junction of the *shha^egfp^* medaka. Magenta letters represent junk insertion by CRISPR-Cas9 editing. (B, C) EGFP expression of *shha^egfp^* medaka in dorsal and anal (B), and caudal fins (C). Dashed lines indicate the outline of fin folds. Standard length (in mm) is given in the upper right of panels B and C. Scale bars: 500 µm (B, C).

**Fig. S9.**
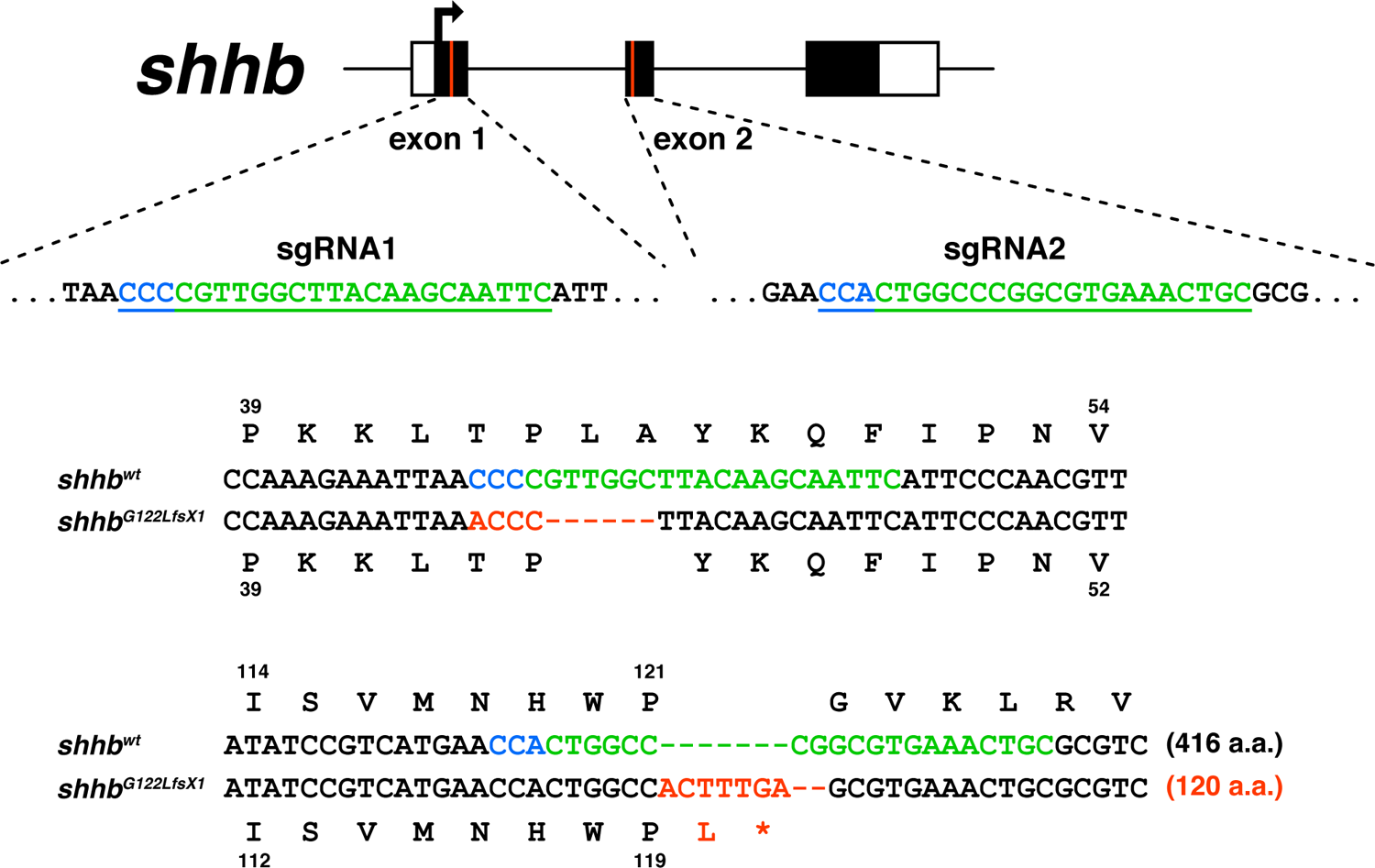
*shhb* mutation in zebrafish. Blue sequences show the protospacer adjacent motif (PAM), green sequences show the recognition sequence for the sgRNA target and red sequences show the mutation site.

**Fig. S10.**
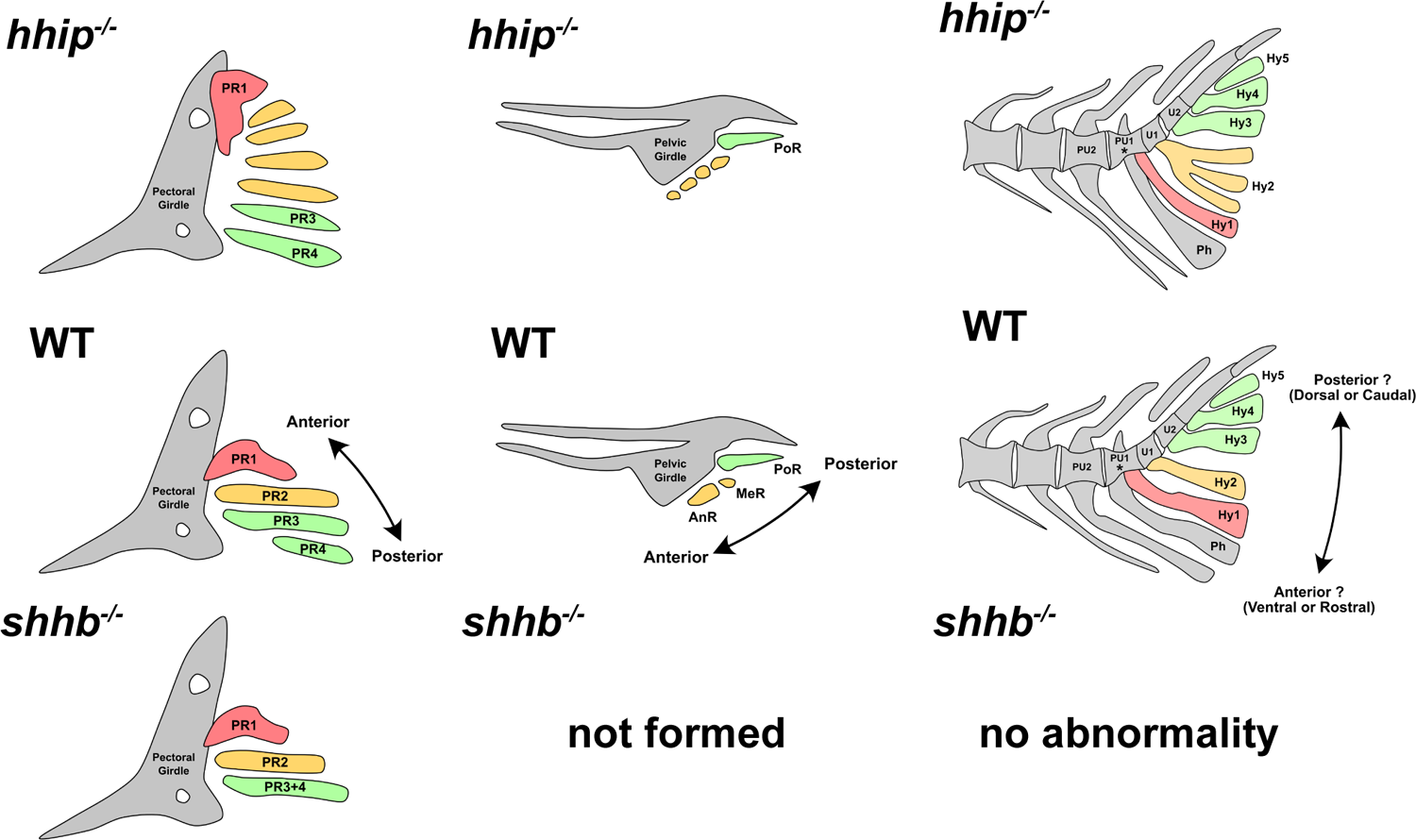
Comparison of the AP identities in paired and caudal fins. Different colors indicate the anterior region (red), the median region (orange), and the posterior region (green).

---

**Table S1.**
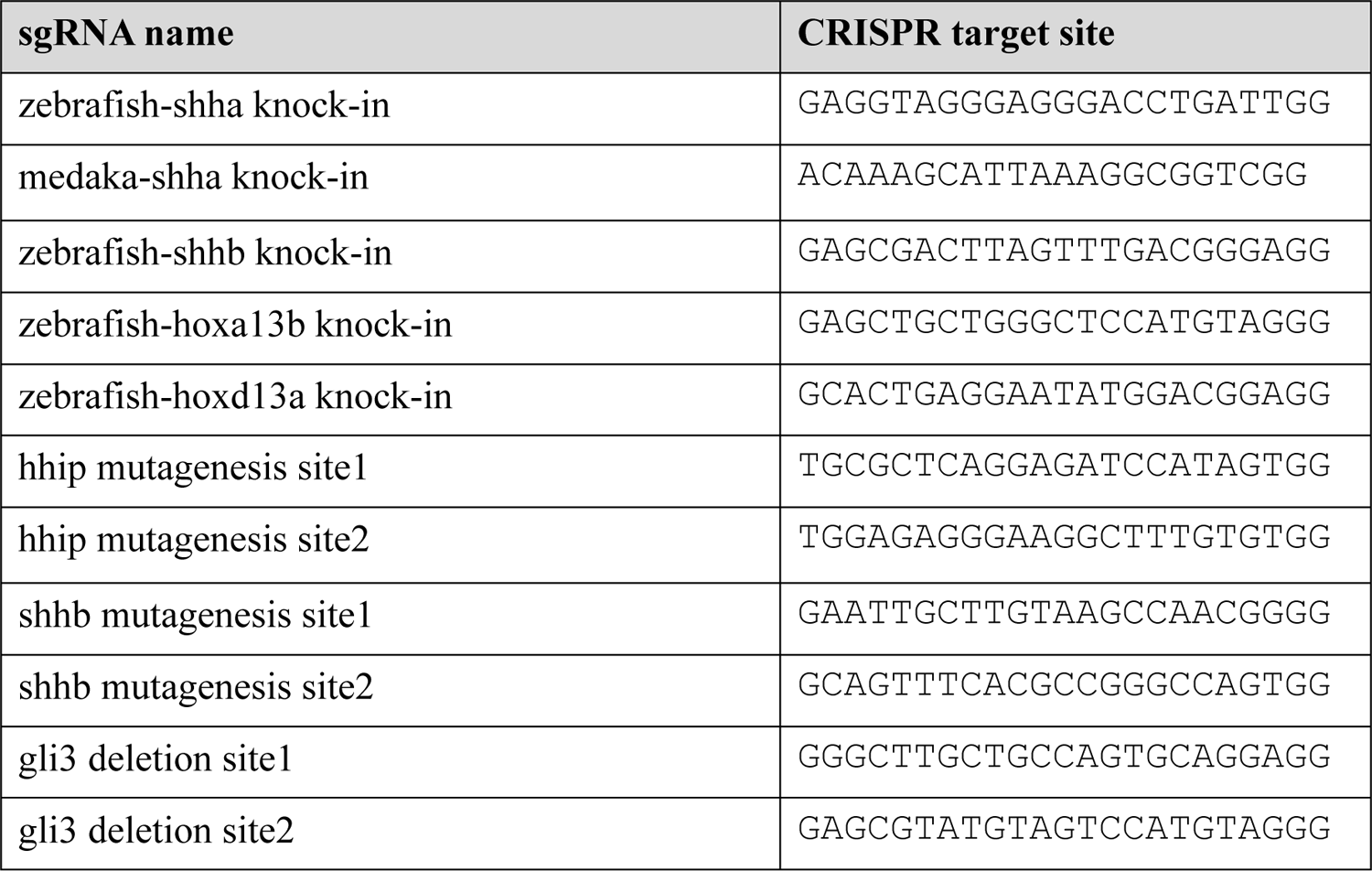
CRISPR target sites.

**Table S2.**
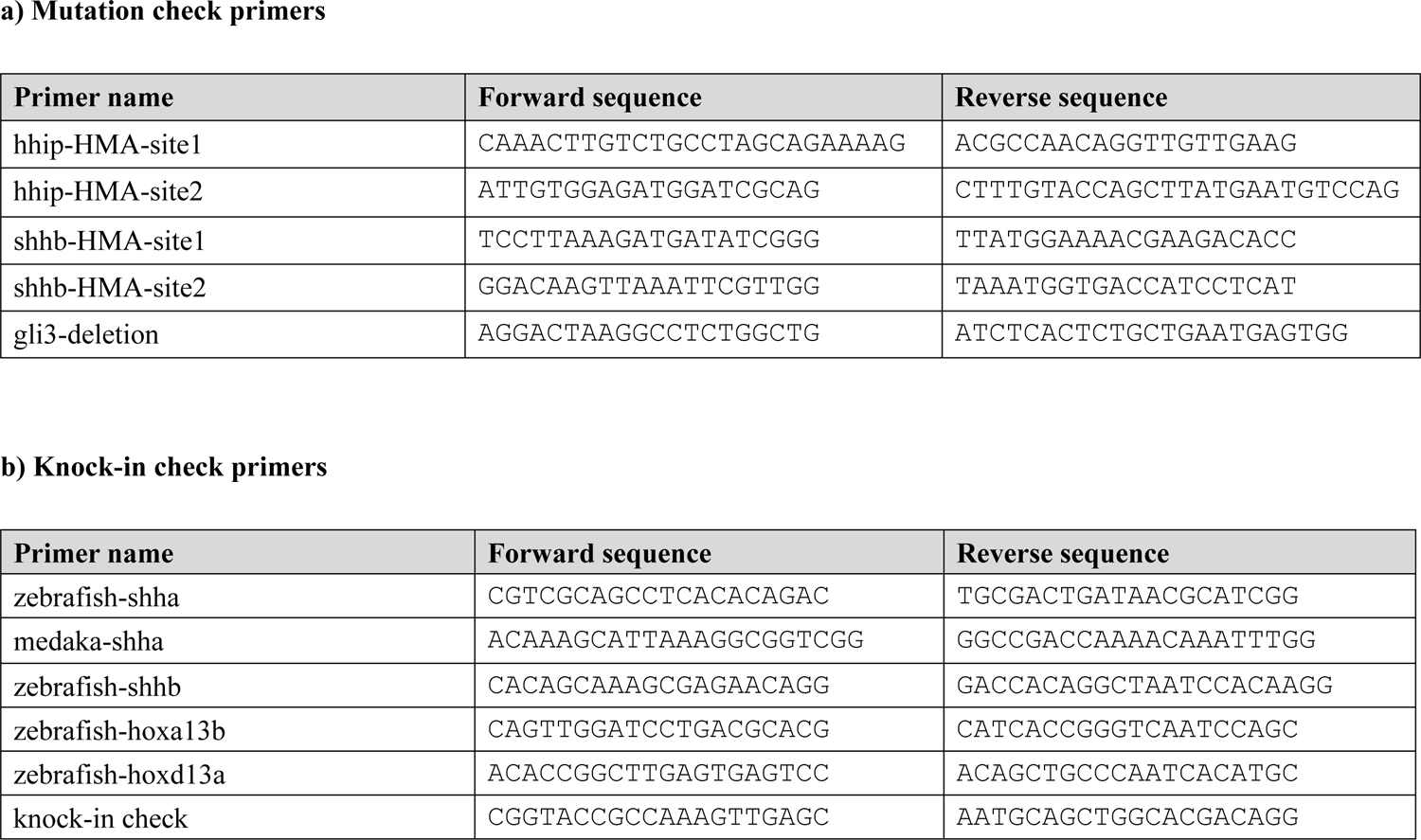
Primer sequences.

**Table S3.**
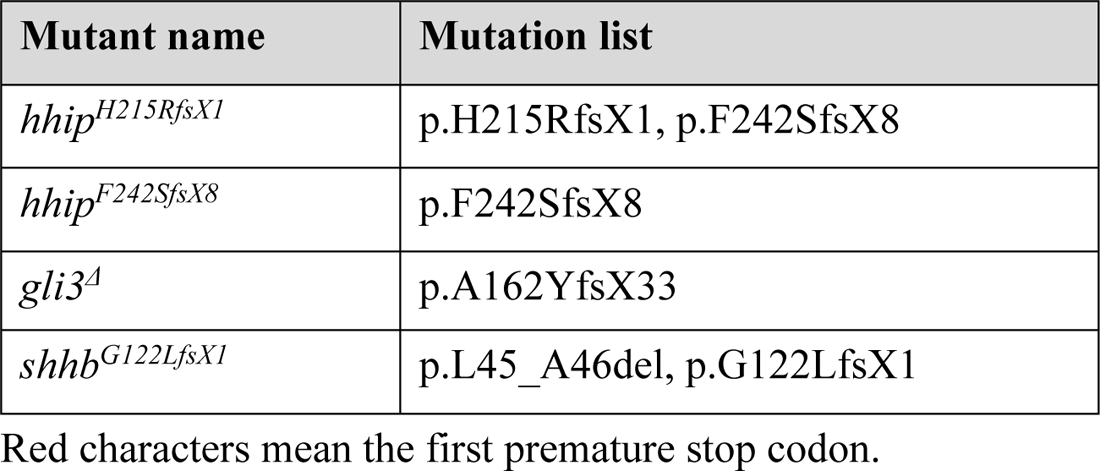
Mutation list.

## Material and Methods

### Fish care and strains

The following transgenic zebrafish lines were used in this study: *col2a1a:EGFP* (51) and *sox10:DsRed* (52). Zebrafish and medaka were housed at 28 °C with light for 14 h, The standard length (SL) of individuals was used as an indicator of individual body size instead of date of development since zebrafish of the same age often have different body sizes (53). All *hhip^-/-^* mutants had compound heterozygous alleles (H215RfsX1/F242SfsX8), due to the high lethality of homozygous alleles (H215RfsX1/H215RfsX1). All *shhb^-/-^*mutants which we observed possessed homozygous alleles (G122LfsX1/G122LfsX1).

### Preparation of sgRNAs

All single guide RNAs (sgRNAs) were designed using CRISPRscan (54) and CHOPCHOP (55) CRISPR online tools. Template DNA for sgRNA synthesize was PCR-amplified using the crRNA/tracrRNA sequence primer, AAAAGCACCGACTCGG-TGCCACTTTTTCAAGTTGATAACGGACTAGCCTTATTTTAACTTGCTATTTCTA-GCTCTAAAAC with the forward primer, AAAAGCACCGACTCGGTGCC, and the reverse primer, TAATACGACTCACTATAggxxxxxxxxxxxxxxxxxxGTTTTAGAGCTA-GAAATAGCA (for T7 polymerase). Lowercase letters indicate genome-targeting sequences (either 19 or 20 mer) in sgRNAs. The genome-targeting sequences in sgRNAs used in this study are shown in Table S1. After PCR amplification with KOD -plus-neo polymerase (Toyobo, Osaka, Japan), PCR products were purified using a PCR purification kit (Cica, Tokyo, Japan). The obtained template DNA was used for in vitro transcription of sgRNAs using a CUGA^®^7 gRNA synthesis Kit (NIPPON GENE, Tokyo, Japan). sgRNAs were purified using a CUGA^®^7 gRNA synthesis kit (NIPPON GENE).

### Microinjection for mutagenesis and knock-in

sgRNAs and Cas9 nuclease were co-injected into one-cell stage zebrafish and medaka embryos. Each embryo was injected with a solution containing ∼10 ng/μl of sgRNA for digesting genomic DNA and ∼250 ng/μl of Cas9 nuclease (IDT, Tokyo, Japan). For knock-in samples, we also added ∼10 ng/μl of sgRNA for digesting BaitD and ∼7.5 ng/μl of phenol-chloroform to extract purified plasmid. To avoid leaky EGFP expression as much as possible, we used pUC-BaitD-Xhbb-EGFP plasmid, which exhibits stable EGFP expression in the target tissues with less leaky expression (33). The injection volume was adjusted to result in the death of approximately 50% of injected embryos within 1 week after injection. These injection mixtures were introduced into one-cell stage eggs, as described previously in zebrafish (56) and medaka (57).

### Genotyping and sequencing of mutant fish

The genotype of mutant fish was determined by heteroduplex mobility assay (HMA) (58). The sequences of mutant alleles isolated from F1 or later generations were determined by direct Sanger sequencing of the PCR products using the primer pair shown in Table S2. All mutations in these mutants are shown in Table S3.

### Genotyping and sequencing of knock-in fish

For knock-in insertion mapping, we collected fluorescent F1 fish at 2-4 dpf and extracted genomic DNA using standard protocols. The insertion status was examined from either the 5′ side or the 3′ side of the insertion. For example, for checking from the 5′ side of the insertion, a PCR reaction was performed using a 5′ primer specific to each target gene (upstream of the expected insertion site) and a 3′ primer specific to the donor plasmid, pUC-BaitD-Xhbb-EGFP. The primers used for insertion mapping in this study are shown in Table S2. The “knock-in check” primers were designed on the 5’ side (fw) and the 3’ side (rv) of the donor plasmid, respectively. The sequences of mutant alleles isolated from F1 or later generations were determined by direct Sanger sequencing of the PCR products.

### Imaging

Zebrafish and medaka were anesthetized by immersion in 0.025% MS222 and placed in 3% agarose gel/E3 on a glass slide. All images were taken using a Leica M205 FA microscopic system and photographed with a Leica DFC 369 FX camera.

### Statistical analysis

The number of skeletal components and the length of pectoral fin primordia were measured at specific time points during the developmental process using *col2a1a:EGFP* fluorescence and Leica Application Suite X (LAS X, Leica). Differences in the number of skeletal elements and the length of between mutant and WT zebrafish were tested using an unpaired t-test using GraphPad Prism (Graphpad Software).

## Acknowledgments

This work was supported by Grants-in-Aid for Scientific Research (KAKENHI) from the Japan Society for the Promotion of Science (JSPS) (KAKENHI Grant number JP20J21314 to Y.T.; JP22K06232 and JP20H04854 to G.A.; JP23H04301 JP22H02627, JP21H05768, JP21K19202 and JP20H05024 to K.T.), and Takeda Science Foundation Life Science Research Grants (Grant number 2022036015 to G.A.).

## Author Contributions

Y.T., G.A., and K.T. designed the research; Y.T., S.O., and S.A. performed the experiments; Y.T. analyzed the data; and Y.T., G.A., and K.T. wrote the paper.

